# Genome-wide predictions of genetic redundancy in *Arabidopsis thaliana*

**DOI:** 10.1101/2020.08.13.250225

**Authors:** Siobhan A. Cusack, Peipei Wang, Bethany M. Moore, Fanrui Meng, Jeffrey K. Conner, Patrick J. Krysan, Melissa D. Lehti-Shiu, Shin-Han Shiu

**Affiliations:** Cell and Molecular Biology Program, Science and Engineering, Michigan State University, East Lansing, MI 48824, USA; Department of Plant Biology, Science and Engineering, Michigan State University, East Lansing, MI 48824, USA; Ecology, Evolutionary Biology, and Behavior Program, Science and Engineering, Michigan State University, East Lansing, MI 48824, USA; Kellogg Biological Station, Michigan State University, East Lansing, MI 48824, USA; Department of Computational Mathematics, Science and Engineering, Michigan State University, East Lansing, MI 48824, USA; Department of Botany, University of Wisconsin-Madison, Madison, WI 53706, USA; Department of Horticulture, University of Wisconsin-Madison, Madison, WI 53706, USA

## Abstract

Genetic redundancy refers to a situation where an individual with a loss-of-function mutation in one gene (single mutant) does not show an apparent phenotype until one or more paralogs are also knocked out (double/higher-order mutant). Previous studies have identified some characteristics common among redundant gene pairs, but a predictive model of genetic redundancy incorporating a wide variety of features has not yet been established. In addition, the relative importance of these characteristics for genetic redundancy remains unclear. Here, we establish machine learning models for predicting whether a gene pair is likely redundant or not in the model plant *Arabidopsis thaliana*. Benchmark gene pairs were classified based on six feature categories: functional annotations, evolutionary conservation including duplication patterns and mechanisms, epigenetic marks, protein properties including post-translational modifications, gene expression, and gene network properties. The definition of redundancy, data transformations, feature subsets, and machine learning algorithms used affected model performance significantly. Among the most important features in predicting gene pairs as redundant were having a paralog(s) from recent duplication events, annotation as a transcription factor, downregulation during stress conditions, and having similar expression patterns under stress conditions. Predictions were then tested using phenotype data withheld from model building and validated using well-characterized, redundant and nonredundant gene pairs. This genetic redundancy model sheds light on characteristics that may contribute to long-term maintenance of paralogs that are seemingly functionally redundant, and will ultimately allow for more targeted generation of functionally informative double mutants, advancing functional genomic studies.

## INTRODUCTION

Genetic redundancy, which refers to multiple genes that perform the same function, has been defined in many ways since the mid-1900s (Gabriel 1960). An early study of genetic redundancy in *Saccharomyces cerevisiae* discussed it in the context of unlinked genes encoding enzymes catalyzing the same reaction (Mortimer 1969). A later study took a broader view of genetic redundancy, with the degree of redundancy ranging from “complete redundancy” among genes with housekeeping functions to “partial overlap of function” among genes with primarily regulatory functions (Pickett and Meeks-Wagner 1995). In studies from a number of model organisms, multiple examples of what is considered genetic redundancy have been given, including: genes derived from convergent evolution encoding enzymes that perform the same function (Pickett and Meeks-Wagner 1995); biochemical pathways that are redundant due to interconnected metabolic networks (Weintraub 1993); and genes from the same family (paralogs) that maintain some of the same functionality (Kempin et al. 1995). Discussions of genetic redundancy in recent literature mostly encompass this last definition, where a duplication event results in multiple copies of a gene that retain overlapping functions (e.g., Chen et al. 2010, Bolle et al. 2013, Rutter et al. 2017). Practically, genetic redundancy is commonly observed as a single gene knockout mutant that shows no phenotype or a mild phenotype compared with a wild-type organism, with a double or higher-order mutant showing a more severe phenotype.

After a gene is duplicated, selection may be relaxed on each copy, allowing accumulation of mutations in one copy, which can lead to pseudogenization (Brookfield 1992); thus, the presence of genetically redundant paralogs long after the duplication event would seem to be an evolutionary paradox (Nowak et al. 1997). In spite of this, the literature is replete with examples of genetic redundancy, and many redundant genes in species such as *S. cerevisiae* and *Caenorhabditis elegans* originated from duplication events that happened over 600 million years ago (Vavouri et al. 2008). At least two mechanisms may explain how this is possible. Redundant copies can be retained for a long time due to the slow pace of genetic drift in large populations. Based on a few key assumptions, it is estimated that a mutation deleterious to the function of a duplicate copy could take 0.75 to 5 million years to be fixed in *Arabidopsis thaliana* (Panchy et al. 2016). However, this cannot account for the apparent redundancy among paralogs from the most recent whole genome duplication (WGD) in the Arabidopsis lineage ~50 million years ago (Bowers et al. 2003). Another possibility is that genetic redundancy is selected for due to its ability to buffer the effect of a deleterious mutation in one paralog (Zhang 2012). The issue is that such a mechanism requires selection based on future needs, which is counter to our understanding of evolution. A mathematical model has been used to demonstrate that redundancy can be stably maintained over time (Nowak et al. 1997). However, the model requirement for perfect equivalency in gene functions and in mutations between paralogs seems unrealistic. Due to the challenges in assessing functions of paralogs, the extent of genetic redundancy and the factors contributing to it remain largely unclear.

Plants are an excellent resource for studying the fate of duplicated genes due to the relatively high rate of WGD events. While pseudogenization (loss of gene function) is the most common fate of duplicated genes in plants (Panchy et al. 2016), some duplicates are retained. By identifying and comparing characteristics of paralogous gene pairs and singleton genes, studies have revealed, for example, a lower synonymous substitution rate among retained (i.e., not pseudogenized) paralogs (Jiang et al. 2013), suggesting that these gene pairs are relatively recent duplicates or that there is selective pressure to retain the ancestral (or a more recently evolved) function. Retention bias is also seen for some gene functions. For example, paralogous transcription factor and signaling genes are retained at a higher rate than DNA repair genes (Blanc and Wolfe 2004). Retention rates of paralogs also vary by duplication mechanism—retained tandem duplicates are more frequently involved in stress responses (Hanada et al. 2008), and genes involved in signaling processes are preferentially retained when derived from WGD rather than smaller duplication events (Maere et al. 2005). While these studies reveal some characteristics of genes that are retained after duplication, they do not directly address whether these retained paralogs maintain redundant functions. A landmark study in Arabidopsis addressed this question using machine learning to integrate 43 gene features related to sequence similarity and gene expression, and predicted that ~50% genes in the Arabidopsis genome have at least one redundant paralog (Chen et al. 2010). In this study, a gene whose single mutant showed no abnormal phenotype (or a mild phenotype) and its closest match in the genome based on sequence similarity were defined as a redundant pair. The most important features for predicting redundancy included differences in isoelectric point, molecular weight, and predicted protein domains between genes in a pair. While this pioneer study provided insights into the prevalence of genetic redundancy, redundancy was defined in only one way without considering the phenotypes of the corresponding double mutants. Also, in the decade since that study substantially more functional genomic data have become available; inclusion of these data in addition to sequence similarity and gene expression may improve the accuracy of redundancy predictions.

While the definition of redundancy presented above is prevalent, observation of unequal genetic redundancy, where the single mutant for one paralog shows a much more severe phenotype than the other and the double mutant has a still more severe phenotype (Briggs et al. 2006), promotes the idea that redundancy is more accurately conceptualized as a continuum. However, the time-consuming nature of precise phenotyping required to quantify redundancy in this manner means that such data are available for relatively few paralogs, and discussions of genetic redundancy frequently exclude single mutants with severe phenotypes. Here we build upon previous work by modeling genetic redundancy using multiple definitions of redundancy by including single mutants in multiple phenotypic categories, and incorporating over 4,000 gene features from six categories, including functional annotations, evolutionary properties, protein sequence properties, gene expression patterns, epigenetic modifications, and network properties. We compared several machine learning algorithms and feature selection methods to identify which of the features have the most predictive power with respect to redundancy. Independent of the model, we additionally performed statistical analysis to identify features common among redundant gene pairs using nonredundant gene pairs as a contrast. To estimate the prevalence of genetic redundancy throughout the genome, we used two of the best-performing genetic redundancy definitions to predict whether ~18,000 gene pairs in the Arabidopsis genome are genetically redundant. Finally, to assess the accuracy of our model, we validated predictions using a “holdout” testing dataset and a handful of experimentally well-characterized gene pairs.

## RESULTS and DISCUSSION

### Definitions of genetic redundancy

The designation of a gene pair as genetically redundant requires phenotype data for double mutants and the corresponding single mutants. To define a set of benchmark redundant and nonredundant gene pairs, we used phenotype data for 2,400 single and 347 higher-order Arabidopsis mutants (including 271 double mutants) from a previous study (Lloyd and Meinke 2012) in which mutants were classified as having no phenotype, a less severe phenotype (i.e., conditional, cellular/biochemical, or morphological), or a severe phenotype (i.e., lethal, indicating the gene is essential) based on comparison with wild-type individuals. We assigned these categories phenotype class numbers: 0 (no phenotype), 1 (conditional), 2 (cellular/biochemical), 3 (morphological), and 4 (lethal) (Figure 1A) and applied this same phenotype classification to 29 additional gene pairs (Bolle et al. 2013), resulting in a final benchmark set of 300 single and double mutant trios (two single mutants and one corresponding double mutant). Note that our data are from experiments generally not designed to assess genetic redundancy and typically conducted in one or a limited number of conditions and environments. Thus, it more straightforward to identify an abnormal phenotype (indicative of nonredundancy) than to prove the absolute absence of an abnormal phenotype (indicative of redundancy).

**Fig. 1.**
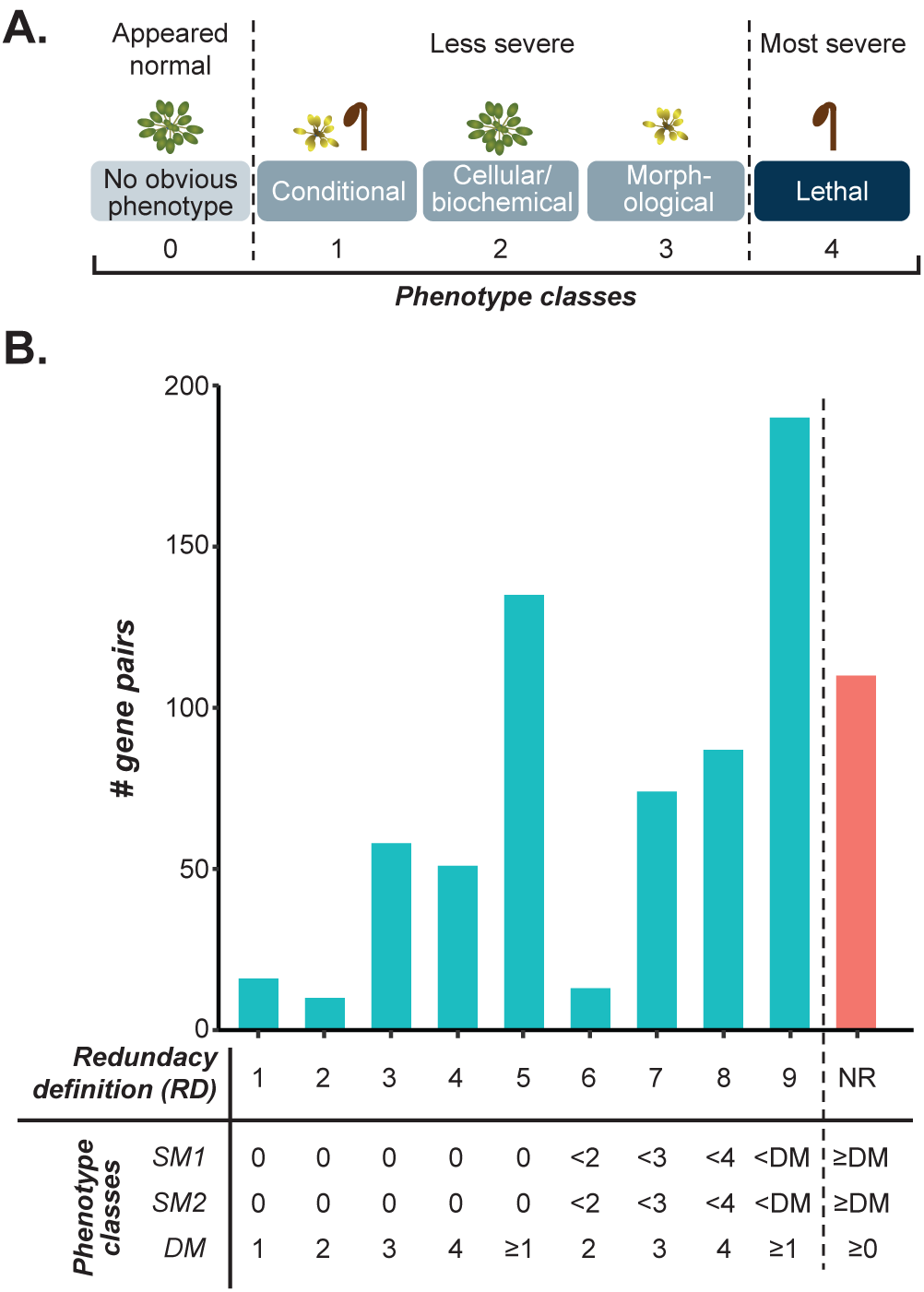
(*A*) Phenotype classes from Lloyd and Meinke 2012. (*b*) Definitions of redundancy and nonredundancy (NR) based on phenotype classes of both single mutants (SM1 and SM2) and the double mutant (DM) for each gene pair. The number of gene pairs assigned to each definition is shown. The comparison signs (≥ and <) are in reference to phenotype category numbers; for example, a phenotype class of “<4” could be 0, 1, 2, or 3. RD5 is RD1-4 combined and RD9 is RD1-8 combined.

Using the benchmark phenotype data and the core idea for defining genetic redundancy based on phenotype severity of single mutants only in comparison to the corresponding double mutant, we established nine redundancy definitions (RD1 to 9) (Figure 1B). The most inclusive definition of genetic redundancy was RD9. Under this definition, a gene pair was considered redundant if the phenotypes of both single mutants were less severe (i.e., assigned a lower phenotype class number) compared with that of the double mutant; 190 benchmark gene trios met this definition. The other RDs were subsets of RD9. Using these more stringent definitions, only mutants of a particular phenotype class were included in the benchmark dataset; for example, when using RD4 only single mutants of class 0 (no phenotype) and double mutants of class 4 (lethal) were considered. The use of multiple definitions offered insulation against errors due to the inherent challenges of classifying phenotypes into specific categories (e.g., some morphological phenotypes are much more severe than others; under specific conditions, conditional lethal is effectively the same as lethal). For example, while RD4 excluded conditional lethal double mutants, as these belonged to phenotype class 1, both types of mutants were included in RD5 and RD9. While we acknowledge that this classification of phenotype severity has caveats, in the absence of quantitative phenotype data on a large scale, quantitative categories together with our multiple definitions of redundancy allow us to better utilize the dataset and begin addressing redundancy more as a continuum than as a binary problem.

To define nonredundant gene pairs, a single definition was used: two genes were considered nonredundant if the double mutant was in the same phenotype class as either single mutant or in a class with a lower number; that is, at least one single mutant had an equal or more severe phenotype than the double mutant (Figure 1B). The nonredundant set contained 110 gene trios. The nearly 2:1 ratio of redundant to nonredundant gene pairs may reflect a bias in the reporting of double mutant phenotypes; a positive result supporting the presence of a more severe phenotype in the double mutant would tend to be reported, with negative results less likely to appear in the literature. Because comparably fewer gene pairs for which the double mutant has no abnormal phenotype have been reported, nonredundant gene pairs very likely are less common in our dataset than they are in nature. Double mutants with much more dramatic phenotypes compared with the single mutants were also overrepresented in our dataset (Figure S1), likely for similar reasons. As a result, some definitions that included only double mutants with mild or no phenotypes had too few gene pairs (RDs 1, 2, and 6, which had 16, 10, and 13 gene pairs, respectively) to generate robust models and were therefore excluded from further analyses.

### Optimal parameters for prediction of genetic redundancy with machine learning

Machine learning allows integration of multiple data types to build a statistical model that can predict a specific outcome. In our case, we were interested in establishing a machine learning model that could predict whether a gene pair was redundant or not using six broad categories of data: functional annotations, evolutionary properties, protein properties, gene expression patterns, epigenetic modifications, and network properties (**Table S1**). The general approach we took is illustrated in Figure 2A. Here the input for the model consisted of benchmark gene pairs (instances), classified as redundant or nonredundant (labels) according to our nine definitions, and information about the genes and gene pairs from the six categories of data (referred to as features). To alleviate the possibility of overfitting our model due to the large number of features examined (~4,000) compared with the number of instances (161-300 depending on the definition), we used 90% of the benchmark gene pairs (training set) to train a model for predicting if a gene pair is redundant or not in a 10-fold cross-validation scheme. Performance was measured using the Area Under the Curve-Receiver Operating Characteristic (AUC-ROC); higher scores indicate a higher true positive rate (proportion of all redundant gene pairs correctly predicted) over the range of false positive rates (proportion of gene pairs incorrectly predicted as redundant). Performance was additionally measured using the Area Under the Precision Recall Curve (AU-PRC); higher scores here indicate greater precision (proportion of gene pair predictions that are correct) over the range of true positive rates ("recall"). Because we used a binary classification scheme (redundant or not) for machine learning, a model classifying gene pairs at random would have a score of 0.5 for both the AUC-ROC and AU-PRC measures, while a perfect model would have a score of 1. Comparing three commonly used machine learning algorithms, Support Vector Machine (SVM), Random Forest, and Gradient Boosting, we found SVM was on average the best-performing algorithm when using RD9 (Figure S2A–B; ANOVA, *p*-value < 2 × 10^−16^, and Tukey’s Honestly Significant Difference test, *p-*values < 0.008).

**Fig. 2.**
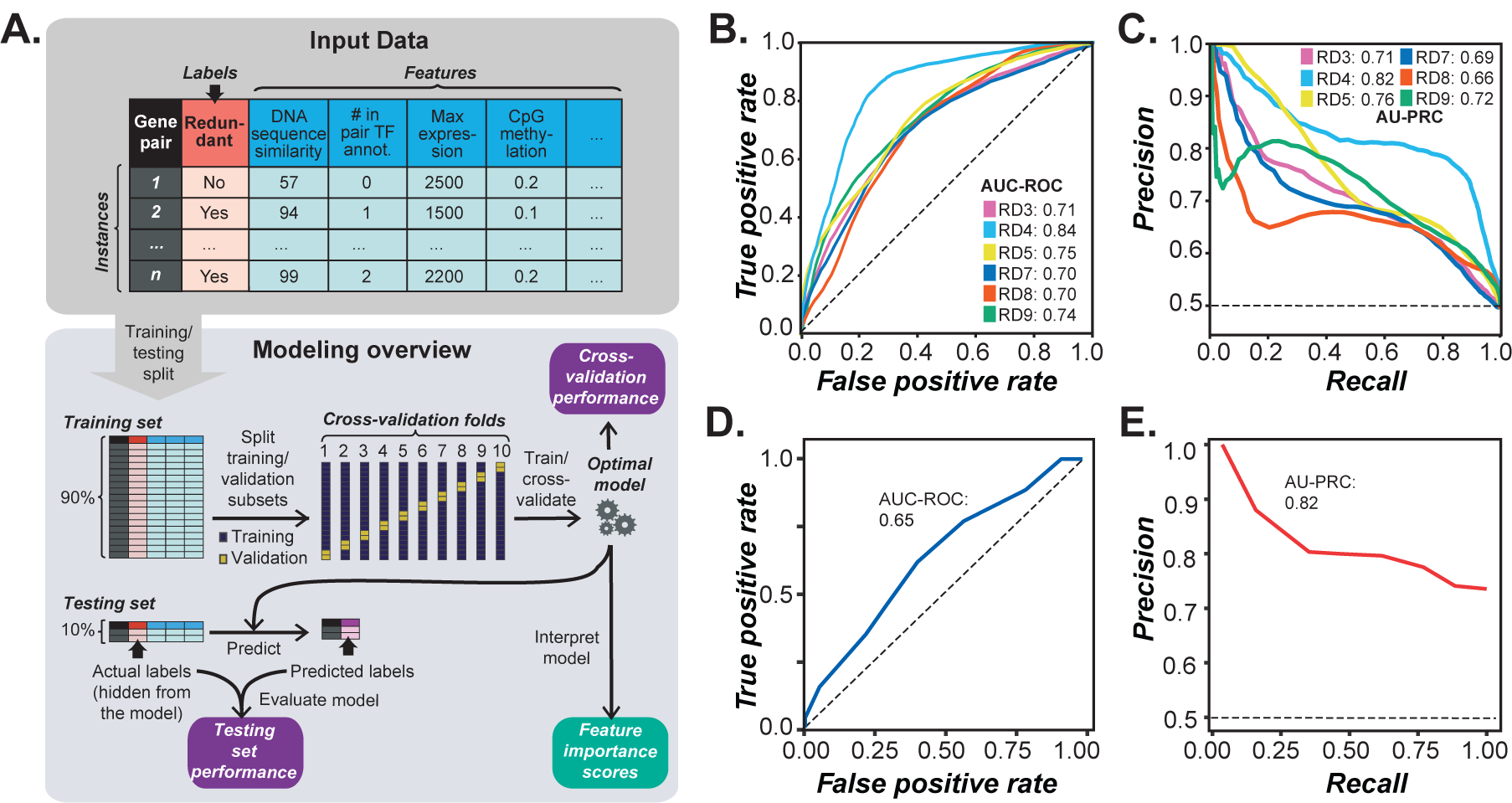
(*A*) Machine learning pipeline workflow. Input data consisted of instances (gene pairs) with labels (redundant or nonredundant) and values of features (characteristics of gene pairs). Example features, as shown in the table, include DNA sequence similarity, the number of genes in a pair annotated as having transcription factor (TF) activity, maximum gene expression level, and the average level of CpG methylation among genes in the pair. The full input data are provided in **Supplemental Data**. Instances were first split into training and testing sets. The training set was further split into a training subset (90%) and validation subset (10%) in a 10-fold cross validation scheme. The optimal model after tuning the model parameters was used to provide performance metrics based on cross-validation, predict labels in the training set for model evaluation purposes, and to obtain feature importance scores. (*B-C*) Cross-validation performance of models built using six of nine RDs based on (*b*) Area Under the Curve - Receiver Operating Characteristic (AUC-ROC) and (*C*) Area Under the Precision-Recall Curve (AU-PRC). RD1, 2, and 6 were not included due to small training data sizes. A model classifying gene pairs perfectly would have AUC-ROC and AU-PRC scores of 1.0; black dotted lines represent the performance of a model classifying at random, in which AUC-ROC and AU-PRC scores would be 0.5 given that we used balanced data (i.e., equal number of redundant and nonredundant instances). (*D*) AUC-ROC and (*E*) AU-PRC for a model trained using RD4 gene pairs and half of the nonredundant pairs (randomly selected) then applied to RD9 gene pairs (excluding RD4) and nonredundant pairs that did not overlap with those used in training the RD4 model.

We next explored how the number of features examined and feature value transformation affected model performance. While models using multiple features generally perform better than those based on single features, the presence of uninformative features can decrease model performance. We tested two methods, Random Forest and Elastic Net, for selecting the most informative features. We looked at the effect of transformation because transforming feature values (e.g., taking the square of values) can amplify small differences, allowing subtle patterns to be more readily identified. We transformed the features four different ways (log, square, reciprocal, and binned) and compared model performance using the original, non-transformed features; the best transformation for each feature (as determined by feature importance scores from the trained models); or multiple transformations of the same original feature. We tested 24 feature combinations (see **Table S2**) by asking how well the model based on each feature combination performed in predicting the RD9/nonredundant benchmark genes in cross-validation. We found that using 200 features selected with Random Forest, using the best transformations of each, led to the best performing model (AUC-ROC = 0.74, Figure S2C and AU-PRC = 0.72, Figure S2D), with a 15% and 18% improvement in performance over a model using all of the untransformed features (AUC-ROC = 0.64, Figure S2E and AU-PRC = 0.61, Figure S2F). The selected features included many that were different representations of the same, raw feature. For example, several features related to total synonymous substitution rate (*Ks*), namely maximum *Ks*, minimum *Ks*, average *Ks*, difference in *Ks*, and total (sum) *Ks* for genes in a pair (see **Methods**) were all among the features selected for RD9, demonstrating that representing a characteristic such as *Ks* in a variety of ways provides distinct and useful information for building the model.

### Comparison of models built with different redundancy definitions

We anticipated that the training sets established using some RDs would result in more accurate predictions than others. Therefore, we next identified the RD that resulted in the best predictions of redundancy using the optimal algorithm (SVM) and input feature set that we identified (200 features selected with Random Forest, using only the best transformation of each feature). When comparing how well each model performed on the cross-validation sets, the model built for RD4 (referred to as the RD4 model) had the best performance (AUC-ROC = 0.84, Figure 2B; AU-PRC = 0.82, Figure 2C; light blue lines). This RD had the highest contrast between single and double mutant phenotype classes (0—no apparent phenotype— and 4—lethal, respectively). A likely reason for the better performance of the RD4 model is that it was easier to build a model to distinguish between redundant and nonredundant gene pairs when the phenotype differences were the most extreme. The second-best models were the ones with the largest training sample sizes, RD5 and RD9 (yellow and green lines, respectively, Figure 2B–C). Thus, it appears that phenotype class contrast and sample size were the most important factors influencing model performance. We therefore focused on models built with the highest phenotype class contrast (RD4) and the largest sample sizes (RD5 and RD9) for further model building.

While the RD4 model performed the best in cross-validation, the majority of redundant gene pairs in the Arabidopsis genome do not have such a high phenotype class contrast. We therefore tested whether the RD4 model would prove useful in predicting redundancy between gene pairs when there were less extreme phenotype differences between the single and double mutants. The RD4 model was applied to a test set composed of RD9 gene pairs (after removing RD4 pairs) and a random subset of half the nonredundant gene pairs. While the AUC-ROC was only 0.62 (Figure 2D), the high AU-PRC score (0.82, Figure 2E) indicated that, as expected from applying a model built with a more conservative definition of redundancy, this model errs on the side of having a higher number of false negatives rather than false positives. Similarly, the RD5 model was applied to a test set composed of RD9 gene pairs (after removing RD5 pairs) and a random subset of half the nonredundant gene pairs. The performance of this model was significantly worse (AUC-ROC = 0.57, Figure S2G; AU-PRC = 0.59, Figure S2H). Thus, the best-performing models for predicting redundancy among gene pairs with all types of phenotype contrasts were those trained on RD4 and RD9; therefore, these two models were used in the following analyses.

### Important evolutionary features in predicting redundant and nonredundant gene pairs

Because the identification of features that are distinct between redundant and nonredundant gene pairs can provide insights about the biological underpinnings of redundancy, we next assessed whether the distribution of values for each feature among the six feature categories was significantly different between redundant and nonredundant gene pairs based on the RD4 and RD9 definitions (see Materials and Methods). For RD4 and RD9, evolutionary properties had the highest percentage of features statistically associated with redundancy (55% and 53% respectively, multiple testing-adjusted *p-*value (*q) <*0.05; Figure 3A–B), and these features tended to be the most significantly correlated with redundancy (median *q*-value of significant features = 0.0003 and 0.004 respectively; Figure S3A–B). Overall, a shared set of 159 features were significantly associated with redundancy in both RD4 and RD9, and there was a correlation between −log(*q*-values) for each feature in RD4 and RD9 (Spearman’s rank ρ = 0.75, *p* < 2.2 × 10^−16^; Figure 3C). This suggested that some features may be significantly associated with redundancy regardless of definition. However, among the top 200 features selected for building the RD4 and RD9 models, we found that only 33% and 25%, respectively, were significantly associated with redundancy when considered individually (Figure S3C–D), highlighting the utility of considering features jointly using machine learning.

**Fig. 3.**
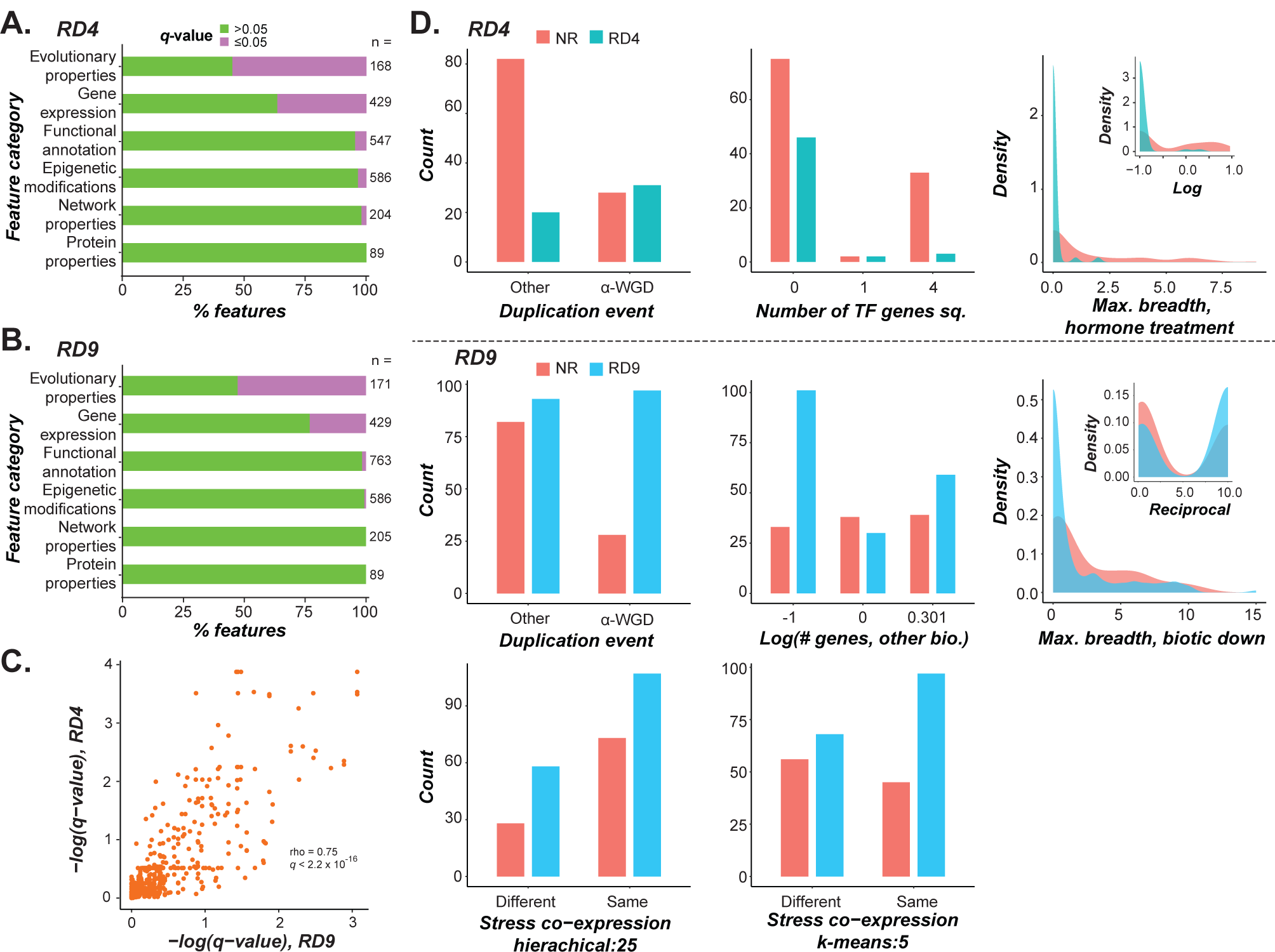
(*A-B*) Percentage of features in each feature category that were significantly associated with redundancy (Wilcoxon rank-sum test for continuous features; Fisher’s exact test for binary features; all multiple-test corrected with Benjamini-Hochberg method) when using (*A*) RD4 and (*b*) RD9. (*C*) Correlation between RD4 and RD9 −log(*q*-values) obtained using the statistical tests as described in (*A*) and (*b*) for each feature. (*D*) Distribution of values among redundant and nonredundant gene pairs for selected features using RD4 and RD9 (separated by a dotted line). For each model, a feature is shown here if the importance score ranked between 1 and 20, was the highest in its feature category, and was significantly associated with redundancy using the statistical tests described in (*A*) and (*b*). For transformed continuous features, untransformed feature values are shown, with transformed values shown as inserts. Abbreviations: “Number of TF genes sq.” is the square of the number of genes in the pair with the annotation DNA-dependent transcription factor; “Max. breadth, hormone treatment” is the maximum number of hormone treatments in which a gene in the pair is differentially expressed. “# genes, other bio.” is the number of genes in a pair with the GO annotation “other biological function”. “Max. breadth, biotic down” is the maximum number of genes in a pair downregulated under biotic stress. “Stress co-expression, hierarchical:25” and “Stress co-expression, k-means:5” refer to co-expression clusters generated from stress datasets with hierarchical (split into 25 clusters) and *k*-means (*k*=5) clustering, respectively; plots indicate whether the genes in a pair are in the same cluster or different clusters.

We next looked into individual features that distinguished redundant gene pairs defined using RD4 and RD9 from nonredundant gene pairs using feature importance scores from the trained models (**Table S3)**. In this case, an importance score represents the degree to which an individual feature contributes to the separation of redundant from nonredundant gene pairs by the algorithm, with features with a higher importance score having a larger contribution. In total, 51 features were shared between the two models (**Table S3**) with well correlated importance ranks (PCC = 0.63, Figure S4A), suggesting that a core set of features are important for predicting redundancy using multiple definitions. However, a shared set of 51 features leaves ~75% of the 200 features selected for each model as unique, highlighting the significant effect of redundancy definition on the models and the types of important features recovered.

The relative importance of the six feature categories ranked from best to worst based on median importance ranks for features in those categories in RD4/RD9-based models was as follows: functional annotations (32/17), evolutionary properties (63.5/81.5), network properties (123/81.5), gene expression patterns (110.5/101.5), epigenetic modifications (108/140), and protein properties (139/133.5). Note that the importance ranks do not mirror the findings in Figure 3A–B, indicating that, for example, while the distributions of >50% of evolutionary property-based features significantly differed between redundant and nonredundant pairs, these features were not as important in predicting redundancy as functional annotation features. The most important feature in both the RD4 and the RD9 models, as determined by feature importance scores, was whether the gene pairs were duplicates from the ɑ-WGD event (for the importance scores of the top 20 features, see Figure S4B–C), with ɑ-WGD-derived gene pairs more likely to be redundant (Figure 3D). The ɑ event is the most recent WGD event in the Arabidopsis lineage, and despite it having likely occurred ~50 million years ago, the importance of this feature suggests that gene pairs derived from this event have not diverged in sequence and function sufficiently to appear nonredundant.

Two other evolutionary property features that were important for both definitions were reciprocal best match (rank=7 and 15 for RD4 and RD9, respectively, Figure S4B–C) and a lethality score-derived feature (discussed below). Reciprocal best matches are paralogous gene pairs that do not have additional retained paralogs generated since their divergence; gene pairs that were reciprocal best matches were more likely to be redundant. As a pair of genes without more recent duplicates are themselves likely to be the product of a relatively recent duplication event (Figure S4D), they are expected to have had less time to diverge in sequence and function, explaining their enrichment among redundant gene pairs. Consistent with this, *Ka* and *Ks*-related features ranked as high as 30 and 32 in the RD4 and RD9 models, respectively. Nonetheless, contrary to our expectations, these evolutionary rate-related features were not the most informative. Instead, other characteristics confounded with rates of evolution, such as mechanism/mode of duplication and, as discussed in the following sections, gene functions and expression profiles, played more important roles in the model.

The reciprocal difference in lethality score was an important feature in both models (rank=2 and 9 for RD4 and 9, respectively, Figure S4B–C). Lethality score is the likelihood that mutation in a gene will lead to a lethal phenotype in Arabidopsis (Lloyd et al. 2015). Thus, we would expect that each gene in a redundant pair would have a low lethality score, and therefore a relatively small difference in lethality score for the gene pair. In contrast to our expectation, we found that redundant gene pairs generally had a smaller difference in reciprocal lethality scores (which equates to a larger difference in raw lethality score) compared with nonredundant gene pairs, although the difference was not significant (Wilcoxon test, *q-*value < 0.11). This unexpected result was likely an artifact of a bias in our data—lethality scores were predicted by Lloyd et al. (2015) for genes without known single mutant phenotypes, but 92% of the genes included in our benchmark dataset have known (nonlethal) phenotypes. In the absence of a predicted lethality score, we used a score of 0 for known nonlethal mutants, which likely artificially lowered the average lethality scores in our benchmark set. Nonetheless, the lethality scores still provided useful information, as indicated by the high importance ranking in both RD4 and RD9.

### Important gene expression, functional, and network features

Features related to gene expression made up the largest portion of features selected for RD4 and RD9 model building, with a total of 126 gene expression features selected for one or both models. The predicted directionality of four features varied between the two definitions, meaning that for a given feature, redundant gene pairs according to one RD had higher values compared with nonredundant gene pairs, while the reverse was true for the other RD. For example, expression variation in the developmental expression dataset (after reciprocal average transformation) was higher for redundant gene pairs according to RD4 than for nonredundant gene pairs, but lower for redundant gene pairs according to RD9. We also found that tissue-specific stress responses varied by redundancy definition; the mean rank of features related to abiotic stress response for RD4 was lower for root tissue (97) than shoot tissue (120), while the opposite was true for RD9 (99 and 94, respectively). Features derived from biotic/abiotic stress and hormone treatment data were more consistently informative across definitions than those from the developmental dataset; while there were four developmental gene expression features in the top 30 for RD9, no such features ranked higher than 54 for RD4. The most important gene expression feature for RD9 was the maximum number of biotic stress conditions under which one or both genes in a pair was downregulated, with redundant gene pairs having a lower maximum than nonredundant gene pairs (Figure 3D). Thus, redundant gene pairs tend not to be downregulated under stress conditions. Previous findings indicated that duplicate genes involved in stress responses are retained at a higher rate than genes involved in other processes (Maere et al. 2005). The most important gene expression feature for RD4 was the maximum number of hormone treatments under which one or both genes in a pair was differentially expressed compared with the control, with nonredundant gene pairs having a higher maximum (Figure 3D).

Among 2,627 functional annotation features, 19 and 13 were among the top 200 for the RD4 and RD9 models, respectively. While only one of these features was selected for both models, given that functional enrichment among redundant gene pairs varies by RD (Figure S5), it was expected that different functional annotation features would be important for predicting redundancy using different redundancy definitions. The most important gene function feature for the RD4 model was the number of genes in a pair (0, 1 or 2) annotated as DNA-dependent transcription factors (referred to as transcription factors). In the trained RD4 model, gene pairs in which both genes had this annotation were more frequently predicted as nonredundant, consistent with the feature value distributions (Figure 3D). This was somewhat unexpected as previous studies have shown that transcription factors are more likely to be retained after gene duplication than other types of genes (Blanc and Wolfe 2004). The most important functional annotation feature for RD9 was the number of genes in the pair having the annotation “other biological processes” (Figure 3D). This term, which encompasses a broad range of processes including responses to stressors or hormones, ion transport, circadian rhythm, aging, and cell growth, among many others, was an important predictor of nonredundant gene pairs.

Finally, while no network properties or protein properties were among the 20 most important features in predicting redundancy for RD4, two network properties were in the top 20 important features for RD9: presence in the same gene co-expression clusters (generated using biotic and abiotic stress datasets, two different clustering algorithms and two different numbers of clusters), with gene pairs in the same cluster more likely to be redundant (Figure 3D). Consistent with this, Chen et al. (2010) found that gene co-expression during pathogen infection was one of the most important features for predicting redundancy in Arabidopsis. In the earlier study by Chen et al. (2010), isoelectric point, overlap in protein domain annotations, and sequence similarity were also among the features found to be important predictors of redundancy. While these features were included in our model building based on RD4 and RD9, they ranked between 26 and 166 depending on the redundancy definition (**Table S3**). Note that the redundancy definition used in the Chen et al. study was based on solely single mutant phenotypes; this likely accounts for the minimal overlap in features found to be important in predicting redundancy.

We also examined the potential causes of mis-predictions by comparing feature values between correctly and incorrectly predicted pairs, generating a score (see Materials and Methods) representing whether mis-predicted nonredundant pairs had feature values similar to RD9 pairs (Figure 4A). We identified several features for which incorrectly predicted nonredundant pairs had values more like RD9 gene pairs than correctly predicted nonredundant pairs, and that also had high feature importance scores, suggesting they may play a role in mis-predictions (Figure 4B). Additionally, in a principal components analysis of correctly and incorrectly predicted nonredundant pairs (Figure 4C), the top 24 features contributing to the first principal component were related to CpG methylation (**Table S4**), implicating it as a major contributor to mis-prediction.

**Fig. 4.**
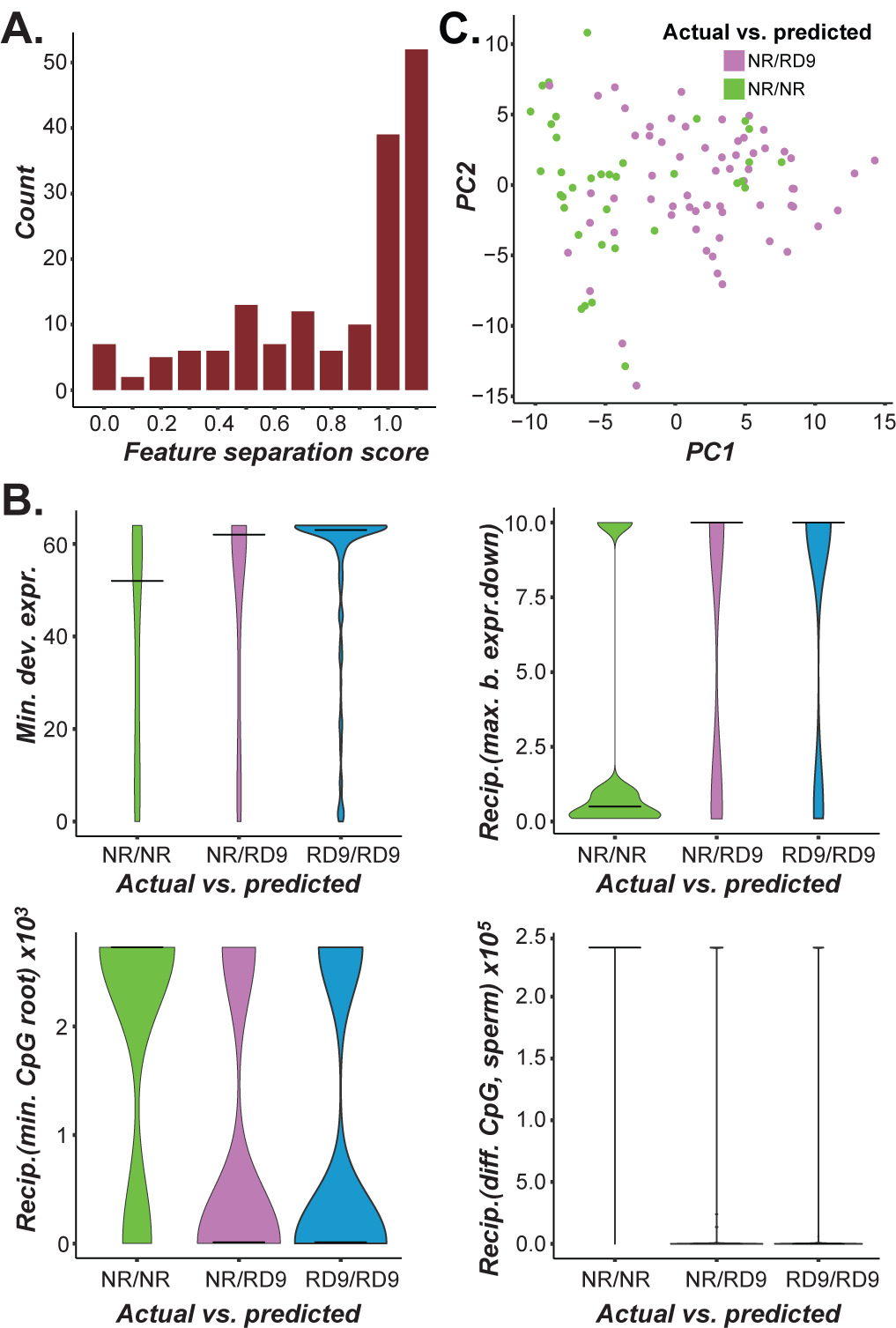
(*A*) Distribution of feature separation scores for features used to build the RD9 model. To identify features that may contribute to mis-predictions, feature values were compared between (1) nonredundant gene pairs predicted as nonredundant (NR/NR), (2) nonredundant pairs predicted as redundant (NR/RD9), and (3) redundant pairs predicted as redundant (RD9/RD9). Using the median value (Med) in each class/predicted class category, we calculated a normalized feature separation score as follows: (*Med*_*NR/RD9*_ - *Med*_*NR/NR*_) /(*Med*_*RD9/RD9*_ - *Med*_*NR/NR*_)For each feature, the feature separation score represents the difference in feature values between correctly and incorrectly predicted nonredundant gene pairs, with a score of 0 meaning that correctly and incorrectly predicted pairs had similar values and a score of 1 meaning that incorrectly predicted pairs had values more similar to redundant gene pairs. Close to 20% of the features had a separation score of 1. (*b*) Distribution of values for selected features among the three categories of actual and predicted redundancy described in (*A*). Horizontal bars indicate the median. “Min. dev. expr.” is the minimum number of tissues and developmental stages in which a gene in the pair is differentially expressed. “Recip. (max. b. expr. down)” is the reciprocal of the maximum number of biotic stress conditions in which one or both genes in the pair are downregulated. “Recip. (min. CpG root)” is the reciprocal of the minimum level of CpG methylation in root tissue for genes in the pair. “Recip. (diff. CpG sperm)” is the reciprocal of the difference in CpG methylation level in sperm cells for genes in the pair. These four features had a feature separation score close to 1 and had feature importance scores in the top 10 for RD9, implicating them in mis-predictions. (*C*) Dimensions 1 and 2 of a principal components analysis performed to identify features that were different between correctly and incorrectly predicted nonredundant pairs. The top 24 features contributing to Dimension 1 were related to CpG methylation levels (**Table S4**).

### Redundancy predictions for Arabidopsis gene pairs not in the benchmark dataset

With the predictive model of redundancy in place, we wanted to get an estimate of genetic redundancy broadly throughout the entire genome. For this analysis, we selected a subset of paralogous gene pairs: all of the WGD and tandem duplicate (TD) pairs in the Arabidopsis genome (7,764 total, collectively referred to as the WG/TD set; **Supplemental Data**). We also sought to model redundancy within a gene family; because a gene family consists of a group of paralogs derived from a variety of duplication mechanisms and with differing evolutionary distances, it offers a wide spectrum of relatedness among gene pairs. For this analysis, we used the protein kinase (Kin) superfamily to generate all possible combinations of gene pairs, then randomly selected 10,000 pairs for analysis (**Supplemental Data**). We expected that applying our model to both datasets would provide information about genetic redundancy at the genome-wide scale and at the more fine-grained gene family level. While both the RD4 and RD9 models showed a high degree of accuracy in predicting redundant gene pairs in cross-validation (87% and 92% of redundant gene pairs correctly predicted, respectively; Figure S6A–B), the RD4 model predicted nonredundant gene pairs with much higher accuracy than the RD9 model (75% and 36%, respectively; Figure S6A–B). Because of the high error rate in predicting nonredundant pairs with the RD9 model, we focused on using the RD4 model to estimate the prevalence of genetic redundancy in the Arabidopsis genome.

Although we analyzed machine learning results primarily as a binary variable (gene pairs were classified as either redundant or nonredundant), these binary predictions were generated from likelihood scores output by the machine learning pipeline. The likelihood score, referred to as a “redundancy score”, ranges on a continuum from 0-1, with 0 being most likely nonredundant and 1 most likely redundant. Using this redundancy score, a threshold score was determined that would maximize the harmonic mean of precision (in this case, the proportion of true redundant pairs to predicted redundant pairs) and recall (proportion of redundant pairs predicted correctly), and this threshold was used to generate the binary predictions for the WG/TD and Kin datasets. Using the RD4 model, the majority of the 17,764 WG/TD and Kin gene pairs were predicted as redundant with redundancy scores well above the threshold (Figure 5). Among the WG/TD set as a whole, 80% were predicted as redundant (Figure 5A), with gene pairs derived from the ɑ-WGD event more likely to be predicted as redundant (83%; Figure 5B) compared with those derived from the β-WGD event (71%; Figure 5C) and the γ and more ancient WGD events (73%, Figure 5D). As duplicate pairs evolve over time, it is expected that the degree of genetic redundancy would continue to decline. While this is true when comparing the ɑ-WGD to older events, similar proportions of duplicate pairs from the β and more ancient events were predicted as redundant based on RD4. This may be because gene pairs derived from the more ancient γ-WGD look similar to those derived from the β-WGD in terms of *Ks* (Maere et al. 2005). However, it is surprising that so many redundant gene pairs (defined based on RD4) that duplicated 50 MYA (ɑ-WGD), 80 MYA (β-WGD; Edger et al. 2015) or longer would be retained. Similarly, 83% of tandem duplicates and 87% of kinases were predicted as redundant based on RD4 (Figure 5E and Figure 5F, respectively).

**Fig. 5.**
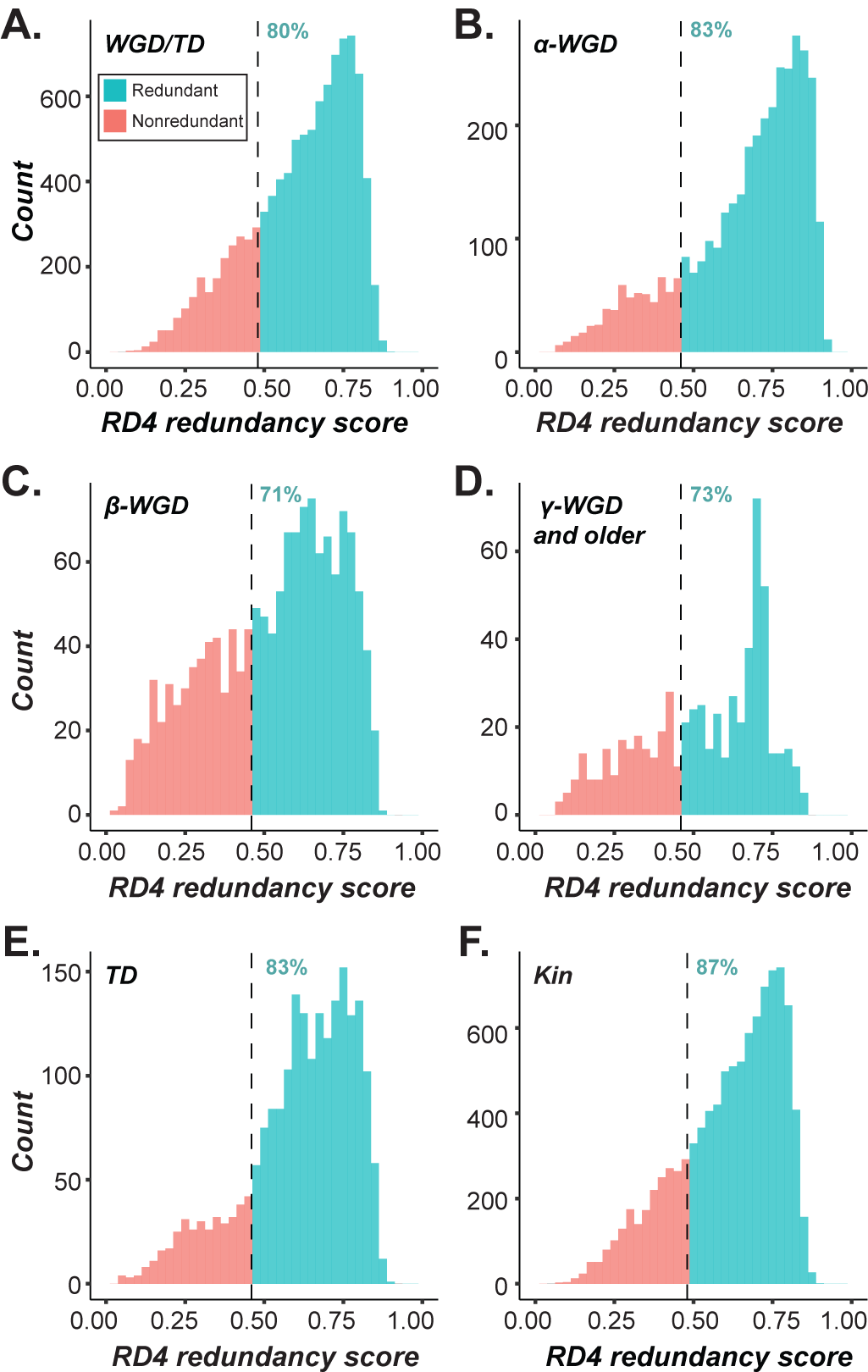
(*A*) Predicted redundancy scores from the RD4 model for gene pairs in the genome derived from whole genome or tandem duplication (WGD and TD, respectively). These results grouped specifically by duplication event/type are shown in the following four sections of this figure. (*b*) Gene pairs derived from the α-WGD event, (*C*) gene pairs derived from the β-WGD event, (*D*) gene pairs derived from the γ-WGD event, (*E*) gene pairs derived from tandem duplication (TD), and (*F*) 10,000 randomly-selected gene pairs from the kinase superfamily (Kin). A majority of gene pairs in all of these datasets were predicted as redundant using RD4.

This percentage of redundant pair predictions was higher than previous estimates in the literature (e.g., Chen et al. 2010). It is important to note that in our WG/TD and Kin datasets, gene pairs are likely being predicted as redundant because they more closely resemble redundant gene pairs with respect to features that have the highest weight in our predictive model (e.g., WGD event). However, the model is built on experimental data that have much more power when calling a gene pair as nonredundant than calling them as redundant; demonstrating that a single mutant has an abnormal phenotype (meaning it is nonredundant) is a simpler task than definitively stating that a mutant has no abnormal phenotype and therefore is redundant with another gene. As previously proposed (Bouché and Bouchez 2001; Bolle et al. 2013), the lack of an observed severe phenotype in a single mutant may be because phenotypes are conditional, tissue-specific, and/or subtle rather than masked by genetic redundancy. Many large-scale phenotyping studies are not able to take these factors into account, and it would therefore be expected that a model built with data from such studies overestimate genetic redundancy in the genome.

While the binary classification of gene pairs as redundant or nonredundant was possible with the available data and straightforward to interpret, it is an over-simplification of the complex nature of genetic redundancy. The threshold-based definition of genetic redundancy may be convenient, but the landscape of genetic redundancy is far more nuanced, with a continuum between gene pairs with various degrees of genetic redundancy. Nonetheless, these data still allowed us to gain valuable insights into the mechanistic underpinnings of genetic redundancy by revealing important features as discussed in the earlier sections. In addition, we anticipate the models can be iteratively improved with the future availability of more phenotype data, particularly quantitative data.

### Validation of predictions

To validate predictions, we used a “holdout” testing set (10%, 16 and 30 pairs for RD4 and RD4, respectively, randomly selected and proportionally divided between redundant and nonredundant pairs, Figure 2A) of the benchmark data. This test set was not included in the model building process and serves to illustrate how the model will perform on new data. Applying the RD4 and the RD9 models on the test set, we obtained AUC-ROC scores of 0.73 and 0.68, respectively (Figure 6A) and AU-PRC scores of 0.62 and 0.82, respectively (Figure 6B). Although there was a decrease in performance compared with cross-validation results (Figure 2B–E), 80% (4/5) and 68% (13/19) of redundant pairs were predicted correctly based on the RD4 and the RD9 models, respectively, and 36% (4/11) of nonredundant pairs were predicted correctly by each of these models (Figure 6C–D). Thus, the holdout testing set generally supported the utility of the RD4 and RD9 models, but the current threshold score was more conservative toward calling gene pairs as non-redundant.

**Fig. 6.**
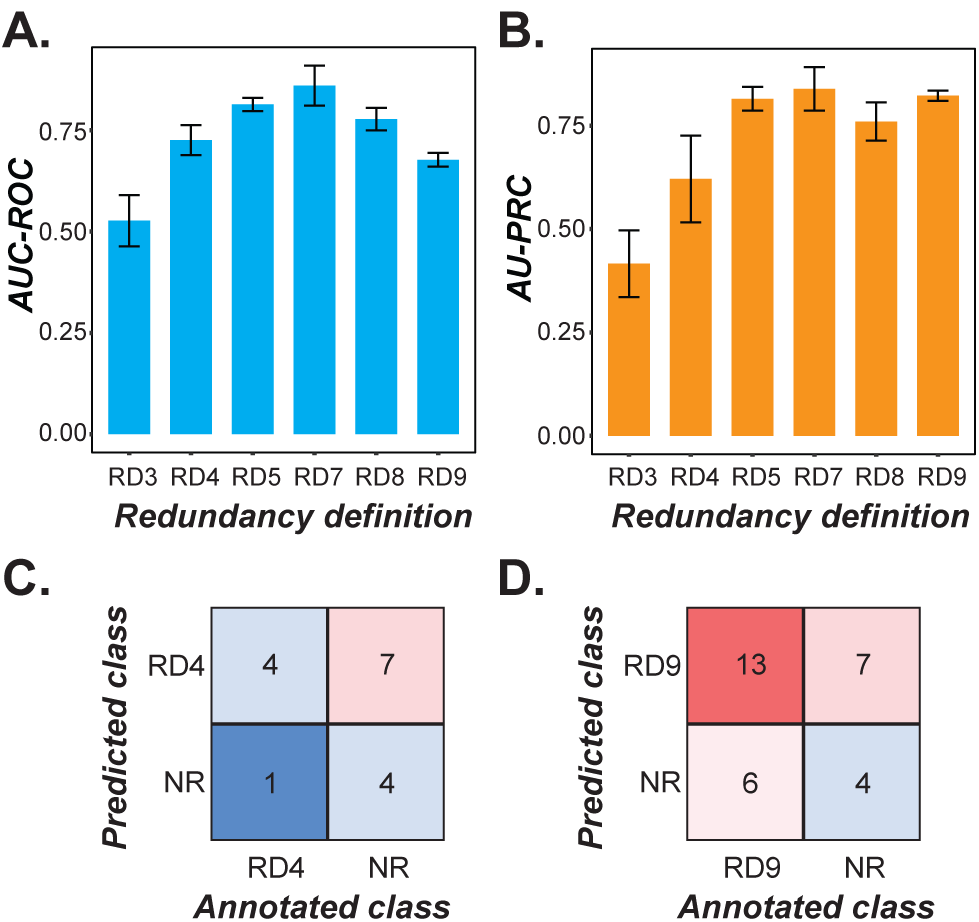
(*A*) AUC-ROC and (*b*) AU-PRC curves for the holdout test sets for models built with each RD. Performance of the models on test sets was lower compared with performance in cross-validation, likely due to the small sample sizes of the test sets. (*C-D*) Confusion matrix for (*C*) RD4 and (*D*) RD9 showing the number of correctly and incorrectly predicted redundant and nonredundant gene pairs in the respective test sets.

Further validation was performed by identifying single and double mutants in the literature that have specifically been studied as mutant trios and have very well documented and characterized phenotypes. We selected ten of these gene pairs: five that meet our criteria for redundancy under RD9 and five we would classify as nonredundant (**Table S5**). Half of the pairs were present in our RD9 benchmark training dataset, while the other half were present in the WG/TD and/or kinase test datasets. We compared the known redundant gene pairs from the literature to our predictions from cross-validation for pairs in the training set and predictions from application of the trained model for pairs in the test set(s). We found that the RD9 model correctly predicted four of five redundant pairs (according to RD9) but mis-predicted all five of the nonredundant pairs as redundant. This comparison was repeated for the RD4 model with the same gene pairs. However, three of the gene pairs classified as redundant using RD9 were classified as nonredundant using RD4 because the double mutants were not lethal. Thus, the validation set for RD4 included two redundant and eight nonredundant gene pairs. The RD4 model correctly predicted one out of the two redundant pairs (according to RD4) and four out of the eight nonredundant pairs. This was consistent with our expectations and prior results showing that the RD9 model tends to err on the side of predicting false positives while the RD4 model is much more conservative and prone to generating false negative predictions.

To determine why mis-predictions may have occurred in these specific cases, we revisited features previously identified as likely contributors to mis-prediction in general in the benchmark dataset (e.g., Figure 4A–B). For the RD9 model, one such feature was reciprocal best match. Although this feature was more strongly associated with nonredundant gene pairs in the benchmark dataset (Figure S7A), the one RD9 pair predicted as nonredundant comprised paralogs that were not reciprocal best matches, making this a likely reason for mis-prediction. Derivation of paralogs from the ɑ-WGD event was another such feature (Figure S7B); three nonredundant pairs predicted as RD9 (nonredundant/RD9) were derived from the ɑ-WGD event, indicating that this feature was a likely contributor to their mis-prediction. Another important feature was related to the number of biotic stress conditions under which genes were downregulated (referred to as biotic downregulation breadth). For this feature, the distribution of feature values among the actual/predicted classes demonstrated that all five nonredundant/RD9 pairs had values more similar to the correctly predicted RD9 pairs than to the correctly predicted nonredundant pairs (Figure S7C). For the RD4 model, the one RD4 pair that was predicted as nonredundant had values for features related to CpG methylation (Figure S7D), gene family size (Figure S7E) and CHH methylation (Figure S7F) that were more similar to those of nonredundant pairs. Additionally, all four of the nonredundant pairs predicted as RD4 had CHH methylation in embryo tissue values that were more similar to those of RD4 gene pairs (Figure S7F).

In total, we identified several types of features that were likely contributors to mispredictions, including duplication event (ɑ-WGD or not), downregulation under biotic stress conditions, and gene methylation patterns. Importantly, we were thus able to identify one or more features that likely contributed to each instance of mis-prediction of both the RD4 and RD9 gene pairs used for validation, a important step in improving future iterations of the model; for example, depending on the definition being used and the importance of the accuracy of predictions (precision) compared with the importance of identifying all redundant gene pairs in a dataset (recall), certain features could be excluded from the model.

### Conclusions

In this study, we optimized and utilized a machine learning approach to predict genetic redundancy among paralogs in Arabidopsis using multiple definitions of redundancy. We identified two biologically relevant and well-performing definitions of redundancy and the optimal 200 features for each definition that allowed us to best model redundancy. Our models performed well on a hold-out testing dataset, demonstrating their utility. Several features related to evolutionary properties, including lethality score, whether genes in a pair were reciprocal best matches, and the type of duplication event from which a gene pair was derived, were consistently ranked as important in generating predictions across redundancy definitions. Interestingly, evolutionary rates, such as *Ka* and *Ks*, were statistically different between redundant and nonredundant gene pairs but not highly ranked in the models, indicating that multiple factors contribute to redundancy, as revealed by machine learning models integrating multiple features. Analysis of these evolutionary-related features demonstrated that redundant gene pairs tend to be more recent duplicates than nonredundant pairs. While it may be tempting to explain redundancy as gene pairs having not had enough time to diverge in function, many redundant pairs are derived from a WGD event estimated to have occurred ~50 million years ago, offering plenty of time for pseudogenization. This suggests that there is some selective pressure to maintain redundancy. In general, we found feature importance to be highly variable by redundancy definition, underscoring the need for testing multiple definitions depending on the biological question being addressed. For example, if one is interested in predicting which genes are lethal or have severe phenotypes a stricter definition is required than when a broader view of redundancy is being used, whereby less extreme phenotype contrasts between single and double mutants would be appropriate.

While the models provide useful information about gene features related to genetic redundancy, there is still room for improvement in terms of prediction accuracy. Performance on test gene pairs withheld from model building was generally not as good as the performance based on cross-validation, which may be due to the small size of the test sets. In addition, our more conservative trained model predicted 84% of 17,764 paralogs throughout the genome to be redundant, which is a much higher estimate than has been shown previously (Chen et al. 2010). This is likely a result of the underlying data used for model building; our models are expected to be biased towards categorization of gene pairs as redundant for the following reasons. We classified redundancy using phenotype data from the literature, including experiments that were not specifically designed to identify redundancy; there are expected to be substantial differences between experiments in how phenotypes were scored. For example, conditional or particularly subtle phenotypes may not have been examined. This likely results in misclassification of single mutants as not having an abnormal phenotype. Because genetic redundancy was defined as a double mutant having more a severe phenotype than the corresponding single mutants, this bias will therefore lead to overestimation of genetic redundancy. Furthermore, classification of gene pairs as redundant or nonredundant, as we were able to do using the broad phenotype categories currently available on a large scale, overly simplifies a complex phenomenon. Redundancy as it exists in nature is not an all-or-nothing binary state, but rather a continuum with a wide range of biologically relevant states.

In our modeling exercise, redundancy scores derived from the model allow an approximation of this continuum, which can be further tested. One approach for testing the degree of genetic redundancy is by obtaining lifetime fitness data for single and double mutant sets. Because lifetime fitness in a mutant reflects the totality of phenotypic effects due to the introduced mutation over the entire life cycle of the individual, subtle and conditional phenotypes are likely better captured. Importantly, our current model can predict redundancy as defined by differences in some phenotypes under some specific conditions. It remains unclear the extent to which such model is relevant to predicting redundancy when it is defined based on single and double mutant fitness, the phenotypic outcome that has the most bearing on the evolutionary fate of a gene pair. Thus, in future studies the generation of lifetime fitness data would allow for a machine learning regression model that more accurately predicts degrees of genetic redundancy between genes in a pair rather than simply classifying genes as redundant or not. Such a model could be applied to gene pairs within a large gene family to compare predicted redundancy scores and reveal patterns related to redundancy maintenance and loss through evolutionary time. Analysis of features important for building the model would be expected to yield additional useful insights about mechanisms related to the evolutionary fate of gene duplicates and the long-term retention of genetic redundancy. Taken together, our results demonstrate the utility of machine learning in combining features to generate accurate predictions of genetic redundancy and identify several evolutionary features that are important in predicting genetic redundancy across several definitions.

## MATERIALS AND METHODS

### Definitions of redundant and nonredundant gene pairs

Arabidopsis mutant phenotype data were collected from Lloyd and Meinke (2012) and Bolle et al. (2013). Our benchmark dataset comprised gene trios for which a double mutant phenotype and both corresponding single mutant phenotypes were reported, with a total of 300 gene trios. A numeric phenotype severity value was assigned to each single and double mutant (Figure 1A), with 0 representing no phenotype; 1, a conditional phenotype of any kind; 2, a cell or biochemical phenotype; 3, a morphological phenotype; and 4, a lethal phenotype. Redundancy was classified using nine definitions (RDs) of varying stringency (Figure 1B). The least stringent definition was RD9, in which any gene pair for which the double mutant phenotype severity score was higher than that of both the single mutants was defined as redundant. With this definition, the dataset contained 190 redundant gene pairs. Gene pairs were classified as nonredundant if at least one single mutant had a phenotype severity score greater than or equal to the double mutant score; the dataset contained 110 nonredundant gene pairs.

### Feature value generation

For predictive modeling, data from six general categories were collected for each gene: functional annotations such as GO terms; evolutionary properties such as synonymous substitution rate; protein sequence properties such as posttranslational modifications; gene expression patterns; epigenetic modifications such as histone methylation; and network properties such as gene interactions (**Table S1**). These data were processed to generate feature values for each gene pair (**Supplemental Data**), and the method used for processing depended on the data type: binary (e.g., whether or not a gene had a given protein domain), categorical (e.g., all the names of protein domains present in a given gene product) and continuous (e.g., gene expression level).

Features such as protein domain and functional annotations were treated as binary and/or categorical input data for feature generation. For processing as binary input data, each gene was assigned a score of 0 (does not have the annotation/property) or 1 (has the annotation/property); gene pair feature values were then generated by taking the number of genes in the pair (0, 1, or 2) having that annotation or property. For example, if Gene1 was annotated as having DNA binding activity but Gene2 was not, the feature value for DNA binding activity for that gene pair would be 1. Additional features were generated by taking the square, −log10, and reciprocal value of features processed in this way. For processing as categorical input data, all annotations of a specific type (e.g., GOslim terms) were listed for each gene. These were then used to represent similarity between genes in a pair. For example, if Gene1 had functional annotations of “DNA binding activity” and “signal transduction” and Gene2 had functional annotations of “signal transduction” and “protein binding”, the number of overlapping annotations would be 1, the total number of unique annotations between the gene pair would be 3, and the percent overlap would be 33. For continuous data, gene pair feature values were generated by calculating the difference, average, maximum, minimum, and total of the values for the gene pair. For example, if Gene1 had an isoelectric point of 10 and Gene2 had an isoelectric point of 9, the difference would be 1, the average 9.5, the maximum 10, the minimum 9, and the total value would be 19. Additional features were generated by taking the square, log10, and reciprocal of features processed as categorical and continuous data, and by assigning each value to one of four quartile bins generated from the untransformed feature data.

### Functional annotation and evolutionary property features

Functional annotations included GO biological process, molecular function and cellular component annotations (The Gene Ontology Consortium et al. 2000; The Gene Ontology Consortium 2017), metabolic pathway annotations from AraCyc v.15 (Mueller et al. 2003), and predicted protein domain annotations from Pfam (Finn et al. 2016). These annotations were processed as binary and categorical data as described above. There were 2,627 features related to functional annotations after transformations were applied (**Table S1** and **Supplemental Data**).

Broadly, evolutionary properties included duplication mechanism and timing, and relationship to other genes in the genome. There were 171 features related to evolutionary properties after transformations were applied (**Table S1** and **Supplemental Data**).

To get the evolutionary rate for each gene in a pair, protein sequences (collected from NCBI; Pruitt et al. 2007) of each *A. thaliana* gene pair were searched against protein sequences from *Theobroma cacao*, *Populus trichocarpa*, *Glycine max* and *Solanum lycopersicum*, using the Basic Local Alignment Search Tool for protein sequences (BLASTP; Altschul et al. 1990). Protein sequences of the gene pair and the best hits in these four species were first aligned using MUSCLE (Edgar 2004), and then were compared to their coding nucleotide sequences to generate the corresponding coding sequence (CDS) alignment. CDS alignments were used to build gene trees using RAxML/8.0.6 (Stamatakis 2014) with parameters: -f a -x 12345 -p 12345 -# 1000 -m PROTGAMMAJTT. *Ka*, *Ks* and the *Ka*/*Ks* ratio on branches leading to each gene of a gene pair were calculated using the free-ratio model of the codeml program in PAML v. 4.9d (Yang 2007). Gene family size and lethality scores were obtained from Lloyd et al. (2015). Where lethality scores were not available, a score of 0 was assigned to known nonlethal genes and 1 was assigned to known lethal genes. Nucleotide and amino acid sequence similarity were calculated using EMBOSS Needle (McWilliam et al. 2013). *Ka*, *Ks*, *Ka*/*Ks*, gene family size, functional likelihood, lethality scores, and sequence similarity were processed as continuous data

Gene pairs were determined to have been derived from one of four types of gene duplication events using MCScanX-transposed (Wang et al. 2013): 1) segmental duplicates— paralogs located in corresponding intra-species collinear blocks; 2) tandem duplicates— paralogs next to each other; 3) proximal duplicates—paralogs close to each other, but separated by ≤ 10 non-homologous genes; 4) transposed duplicates—one of the paralogs located in inter-species collinear blocks, the other not. Segmental duplicates were additionally noted as being derived or not derived from the α- or β-WGD events. Protein sequences of *A. thaliana* were searched against protein sequences of *A. thaliana* (intra-species), *Arabidopsis lyrata*, *Brassica rapa*, *Carica papaya*, *P. trichocarpa*, and *Vitis vinifera* (inter-species) using BLASTP, with a cutoff E-value of 1×10^ℒ10^. Five different sets of parameters were evaluated for MCScanX-transposed: 1) -k 50 -s 5 -m 25; 2) -k 50 -s 2 -m 25; 3) -k 25 -s 2 -m 25; 4) -k 25 -s 2 -m 50; 5) -k 25 -s 5 -m 25; where -k indicates the cutoff score of collinear blocks, -s specifies the number of matched genes required for the calling of a collinear block, and -m means the maximum number of genes allowed for the gap between two genes. The duplication mechanisms inferred using these five different sets of parameters were consistent with one another for the majority of gene pairs; 78 pairs had discrepant results, representing 0.4% of the total dataset. In these cases, the mechanism that occurred most frequently in the results for that gene pair was assigned; if there was no majority, the mechanism was listed as N/A. Each gene pair was assigned a binary value indicating whether or not the genes were reciprocal best matches (i.e., they were one another’s best hit based on nucleotide BLAST searches) and whether or not they were derived from each type of duplication mechanism (e.g., a gene pair derived from the α-WGD event would have a value of 1 for the WGD feature and for the α-WGD feature, and a value of 0 for all other duplication mechanisms).

Retention rate was based on the presence or absence of a paralog in 15 species: *A. lyrata*, *Capsella rubella*, *B. rapa*, *T. cacao*, *P. trichocarpa*, *Medicago truncatula*, *V. vinifera, S. lycopersicum*, *Aquilegia coerulea*, *Oryza sativa*, *Amborella trichopoda*, *Picea abies*, *Selaginella moellendorffii*, *Physcomitrella patens*, and *Marchantia polymorpha*. The retention rate for each gene was calculated as the number of genomes in which a paralog was present divided by the total number of genomes analyzed (16: *A. thaliana* plus the 15 additional species). Genome data were collected from Phytozome (Goodstein et al. 2012) for *P. patens* 318 v3.3, *M. polymorpha* 320 v3.1, *S. moellendorffii* 91 v1.0, *A. trichopoda* 291 v1.0, *O. sativa* 323 v7.0, *B. rapa* 277 v1.3, *C. rubella* 183 v1.0, *A. thaliana* 167 TAIR10, *A. lyrata* v2.1, *M. truncatula* 285 Mt4.0 v1, *V. vinifera* 145 Genoscope 12x, *A. coerulea* v3.1, *P. trichocarpa* 210 v3.0, and *T. cacao* 233 v1.1; from NCBI for *S. lycopersicum* v2.5; and from PlantGenIE (Sundell et al. 2015) for *P. abies* v1.0.

### Gene expression and epigenetic modification features

Processed microarray gene expression datasets were obtained from Moore et al. (2019) and contained gene expression levels under biotic (Wilson et al. 2012) and abiotic stress (Kilian et al. 2007; Wilson et al. 2012), under hormone treatment (Goda et al. 2008), at different developmental stages (Schmid et al. 2005), and at different times of day (Mockler et al. 2007). In addition to these gene expression levels, we also considered expression breadth, which represents the number of tissues and conditions under which each gene is expressed. Gene expression levels and ribosome occupancy from RNA-seq and Ribo-Seq experiments in root tissue were obtained from Hsu et al. (2016) and processed along with the microarray gene expression data as continuous data. There were 450 features related to gene expression after transformations were applied (**Table S1** and **Supplemental Data**).

Epigenetic modifications included DNA methylation, chromatin accessibility, and histone modifications. Percent CHH, CHG, and CpG methylation, gene body methylation, and histone modification data were obtained from Lloyd et al. (2015). Percent methylation values were treated as continuous data, and gene body methylation and histone modification data as binary data. Chromatin accessibility data were from Sullivan et al. (2014) and were also binary, with each gene receiving a score of 1 if it contained a DNase peak site and a score of 0 if it did not. There were 565 features related to epigenetic modifications after transformations were applied (**Table S1** and **Supplemental Data**).

### Protein sequence and network property features

Protein sequence properties included amino acid length, isoelectric point, and posttranslational modifications. Amino acid lengths were obtained from Lloyd et al. (2015). Isoelectric points and myristoylation data were from The Arabidopsis Information Resource (Berardini et al. 2015). Amino acid length and isoelectric point were processed as continuous data. Acetylation, deamination, formylation, hydroxylation, oxidation, and propionylation data were obtained from The Plant Proteome Database (Sun et al. 2009). Posttranslational modifications were processed as binary data: whether or not the protein product was predicted or known to have the modification. In total, 93 features were related to protein sequence properties after transformations were applied (**Table S1** and **Supplemental Data**).

Network properties were related to known or potential interactions of genes or protein products. Gene interaction data (AraNet v.1, Lee et al. 2010) and protein-protein interactions (AtPIN, Brandão et al. 2009) were processed as categorical data. Gene co-expression was calculated from the microarray datasets referenced above using multiple clustering algorithms, namely k-means, c-means and hierarchical clustering at k=5, 10, 25, 50, 100, 200, 300, 400, 500, 1000, and 2000 as described in Moore et al. (2019). These data were processed as categorical data, with each combination of clustering algorithm, dataset and k-value included as a feature; a gene pair received a value of 1 if both genes were in the same cluster and a value of 0 if they were not. There were 205 features related to network properties after transformations were applied (**Table S1** and **Supplemental Data**).

### Identification of features distinguishing redundant and nonredundant pairs

To identify features that could distinguish between gene pairs from the redundant and nonredundant classes, we applied statistical tests to determine if feature values were significantly different between the classes. Binary gene pair features (e.g., duplication type, presence in a gene co-expression cluster) were analyzed using two-sided Fisher’s exact tests with multiple testing correction using the Benjamini-Hochberg method (Benjamini and Hochberg 1995). To determine whether feature value transformations improved the ability to distinguish between classes, the reciprocal, square, and log10 of continuous features were included as separate features. Continuous values were also binned into four quartiles of equal size and bin values included as features. Transformed and untransformed continuous feature values were analyzed using a Wilcoxon rank sum test (Wilcoxon 1945) with multiple testing correction performed using the Benjamini-Hochberg method. Features were considered to distinguish between redundant and nonredundant gene pairs if *q* < 0.05 after multiple testing correction (**Table S1**). Continuous feature effect sizes are the standardized z statistic (calculated from the *p*-values given by the Wilcoxon rank sum test) divided by the square root of the sample size. Binary feature effect sizes correspond to the odds ratio calculated from the enrichment table for each feature.

### Redundancy prediction model building and optimization with machine learning

Models for predicting genetic redundancy between gene pairs were built with Random Forest, Gradient Boosting and SVM algorithms implemented in the scikit-learn machine learning package (Pedregosa et al. 2011) in Python. For Random Forest and Gradient Boosting, a grid search was performed with 10-fold cross-validation for parameter optimization; gene pairs were randomly divided into a training set (90%) and a testing set (10%) with proportional division of redundant and nonredundant gene pairs. This division was repeated 10 times, with 10 replicate models built for each iteration, for a total of 100 models. Ten-fold cross-validation was also used in model building with 100 iterations each for a total of 1000 models, with each being tested on a “holdout” testing dataset, consisting of 10% of the benchmark dataset and not included in building the model, to assess performance. The trained model was used to predict redundancy among all tandem and WGD pairs in Arabidopsis (**Supplemental Data**) and among a random sample of Arabidopsis kinase gene pairs. Using kinase family classifications from Lehti-Shiu and Shiu (2012), all possible within-family combinations of gene pairs were generated. Ten thousand of these pairs were then randomly selected for predictions (**Supplemental Data**).

## DATA AVAILABILTY

**Supplemental data** will be available at http://zenodo.org.

## ACKNOWLEDGEMENTS

We thank Christina B. Azodi for assistance with machine learning methods and John P. Lloyd for providing processed data. This work was partly supported by the National Science Foundation (IOS-1546617, DEB-1655386) to J.K.C., P.J.K. and S.-H.S., and U.S. Department of Energy (Great Lakes Bioenergy Research Center BER DE-SC0018409) to S.-H.S.

---

**Fig. S1.**
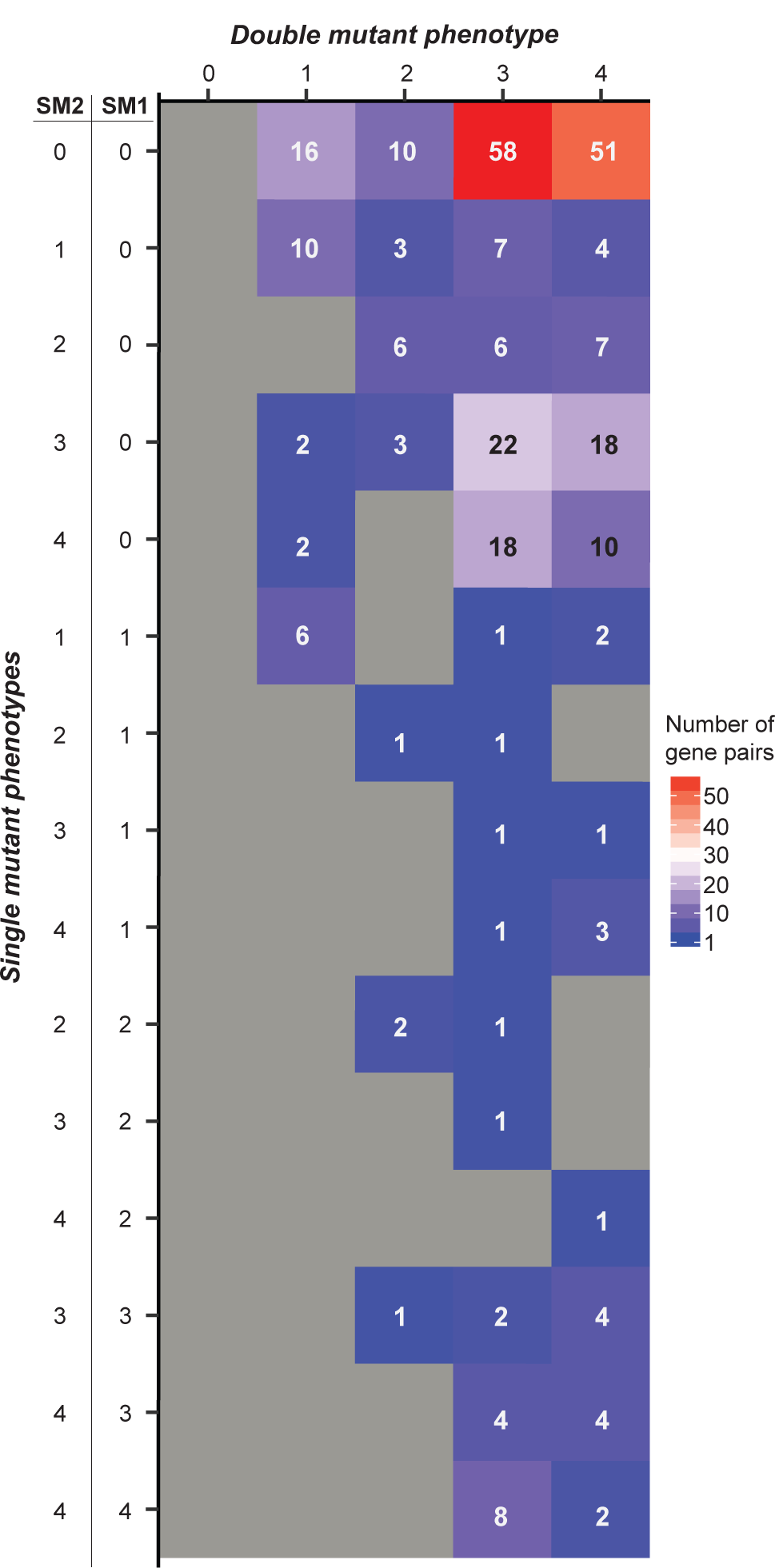
Distribution of benchmark gene pairs among phenotype severity categories (as defined in Fig. 1) for both single mutants (SM1 and SM2) and the double mutant for each pair. The dataset is biased toward double mutants with more severe phenotypes.

**Fig. S2.**
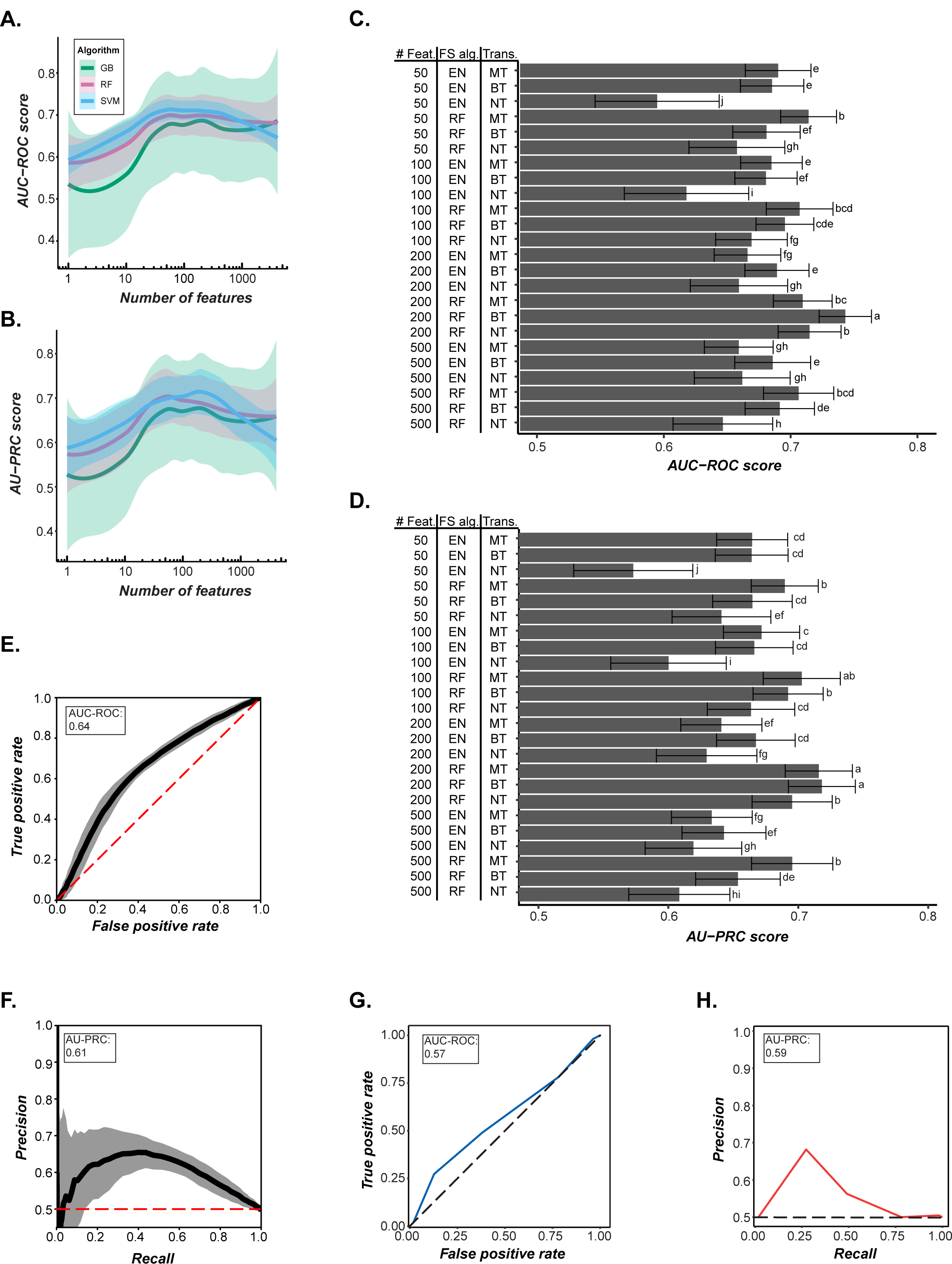
(*A*) AUC-ROC scores and (*B*) AU-PRC scores for binary classification machine learning models built using RD9 with Gradient Boosting (GB), Random Forest (RF) and Support Vector Machine (SVM) algorithms and using different numbers of features. Shading indicates the standard deviation. Using AUC-ROC as a measure, models built with SVM performed the best (ANOVA, *p*-value < 2 × 10^−16^, and Tukey’s Honestly Significant Difference [HSD] test, *p-*values < 0.008). Using AU-PRC, models built with SVM performed significantly better compared with those built with Gradient Boosting (ANOVA, *p* < 2 × 10^−16^; Tukey’s HSD, *p*-value < 0.0001), but not with those built with Random Forest (Tukey’s HSD, *p*-value = 0.36). (*C*) AUC-ROC scores and (*D*) AU-PRC scores for machine learning models built using different combinations of feature numbers (“# Feat.”), feature selection algorithms (“FS alg.”), and numbers of transformations allowed for each feature (“Trans.”). Different letters indicate statistically significant differences between models according to Tukey’s HSD. Using AUC-ROC as a measure, the best-performing combination was 200 features selected with Random Forest and with only the best transformation of each feature allowed (ANOVA, *p* < 2 × 10^−16^; Tukey’s HSD, *p-*values < 0.0001). Using AU-PRC as a measure, this combination was significantly better than all other combinations of parameters (ANOVA, *p* < 2 × 10^−16^; Tukey’s HSD, *p*-values < 2.3 × 10^−4^) except for the following two combinations: 200 features selected with Random Forest, with multiple transformations of each feature allowed, and 100 features selected with Random Forest, with multiple transformations of each feature allowed (Tukey’s HSD, *p*-values 1.00 and 0.13, respectively). (*E*) AUC-ROC curve and (*F*) AU-PRC curve of a model built with all untransformed features, demonstrating the improved performance of the optimized model in (*C*) and (*D*) with respect to both measures. (*G*) AUC-ROC and (*H*) AU-PRC for a model trained using RD5 gene pairs and half of the nonredundant pairs (randomly selected) then applied to RD9 gene pairs (excluding RD5) and nonredundant pairs that did not overlap with those used in training the RD5 model. Using these performance measures, this model did not perform as well as a model trained on RD4 then applied to a test set composed of RD9 gene pairs (after removing RD4 pairs) and a random subset of half the nonredundant gene pairs (Figure 2D–E); therefore, RD5 was not selected for further analysis.

**Fig. S3.**
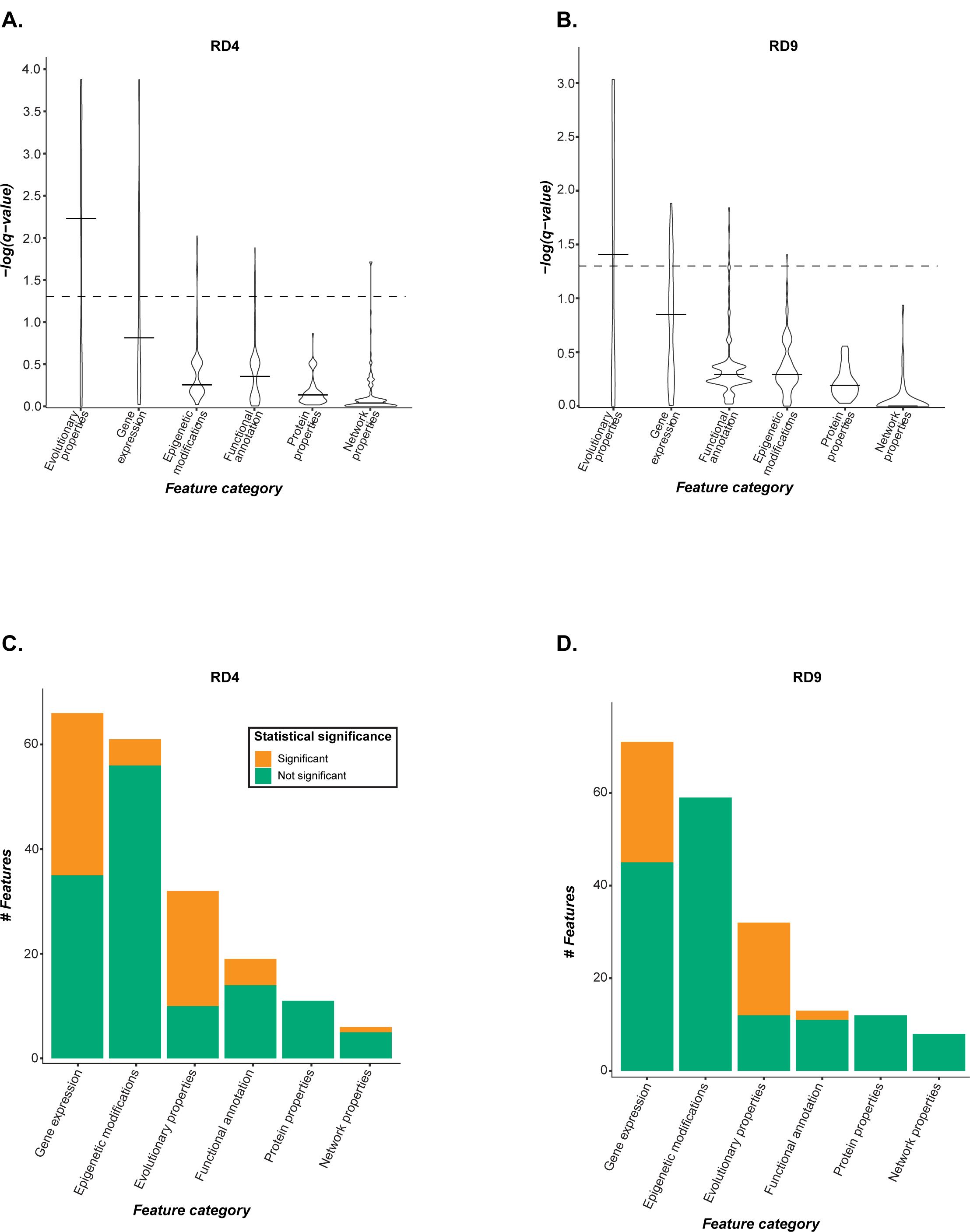
(*A-B*) Distribution of −log(*q*-values) from tests of feature association with redundancy as defined using (*A*) RD4 and (*B*) RD9. Statistical significance was determined with Wilcoxon rank sum test for continuous features and two-sided Fisher’s exact test for binary features; all values were corrected for multiple testing with the Benjamini-Hochberg method. The dotted lines show a *q*-value of 0.05. All *p*-values, *q*-values and effect sizes are reported in **Table S1**. The median effect sizes (calculated as described in **Methods**) were 0.11 (RD4) and 0.7 (RD9) among continuous features, and 1.4 (RD4) and 1.1 (RD9) among binary features. (*C-D*) Distribution by feature category of the 200 features selected for model building for (*C*) RD4 and (*D*) RD9. Features that have a statistically significant association with redundancy as described above are shown in orange. Only 25% of the features selected were significantly associated with redundancy.

**Fig. S4.**
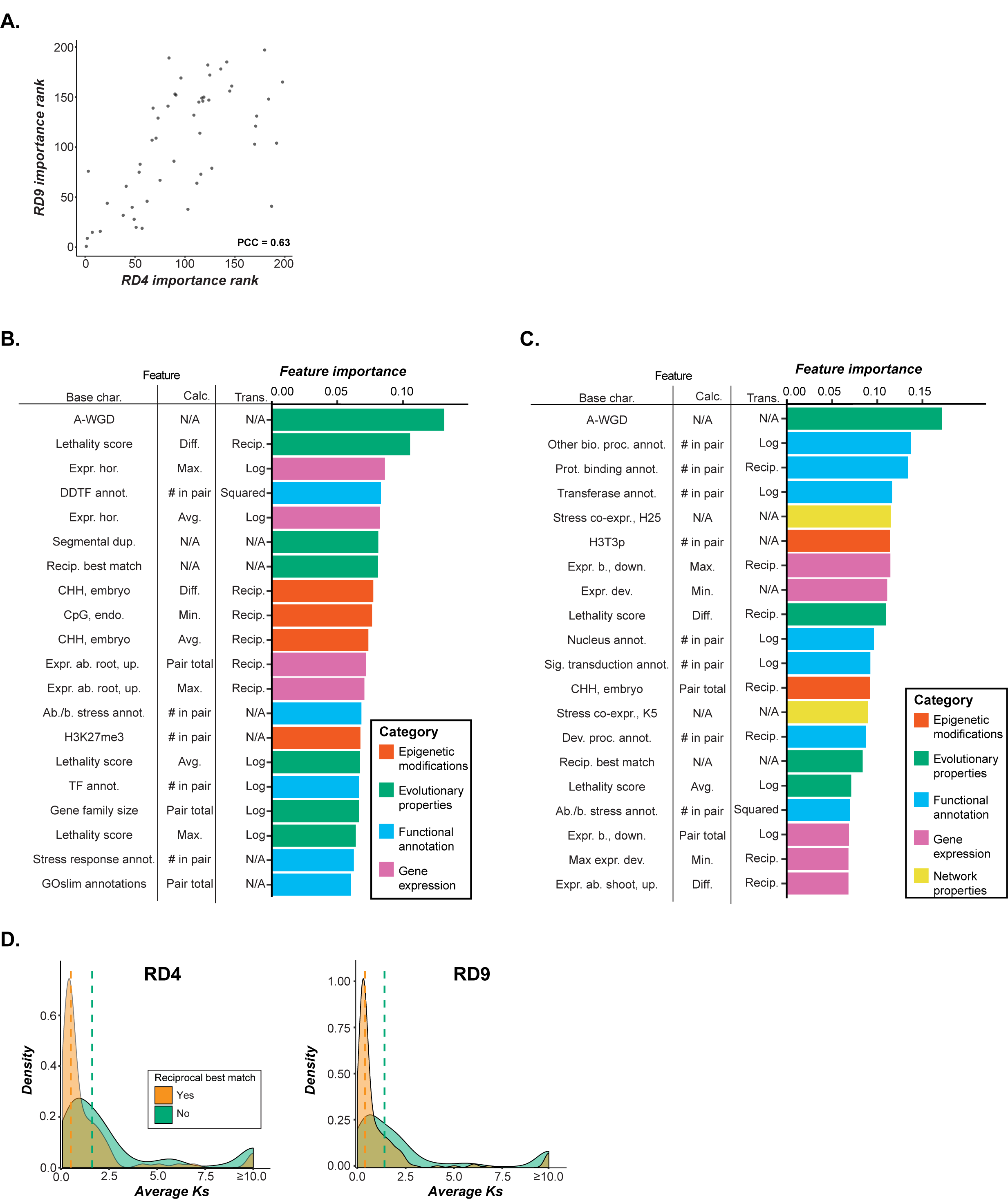
(*A*) Comparison of feature importance ranks of the 51 features included in both the RD4 and RD9 models. The feature importance ranks are well correlated between the two models (PCC = 0.63; *p* = 6.0 × 10^−7^), indicating that a core set of features is important in predicting redundancy across definitions. (*B-C*) Feature importance scores obtained from machine learning models for (*B*) RD4 and (*C*) RD9. (*D*) Distribution of *Ks* values among gene pairs included in the RD4 (left) and RD9 (right) models that are reciprocal best matches (orange) and not reciprocal best matches (green). Dotted lines show the median *Ks* value for each group. Gene pairs that are reciprocal best matches tend to be more recent duplicates as shown by the lower *Ks* values.

**Fig. S5.**
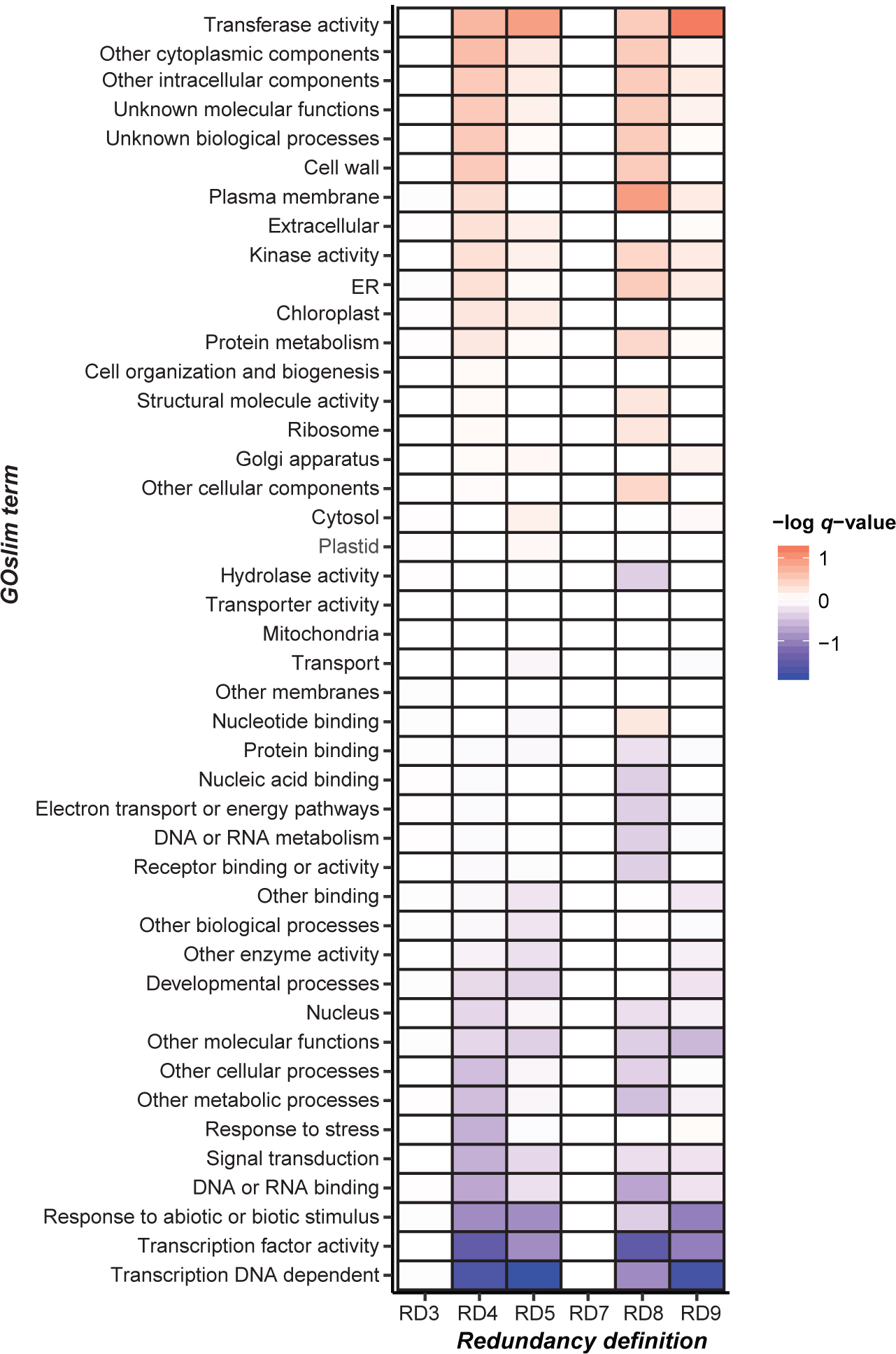
Enrichment of GO terms among redundant gene pairs vs. nonredundant gene pairs for each RD. Blue represents enrichment among nonredundant pairs while red represents enrichment among redundant gene pairs; lighter shades show statistically weaker associations (i.e., higher *q*-value) and darker shades show statistically stronger associations (lower *q*-value). Statistically significant enrichment was seen in transcription factor activity among nonredundant pairs compared with RD4 and RD8 gene pairs, and in DNA-dependent transcription factor activity among nonredundant gene pairs compared with RD4, RD5 and RD9 gene pairs. In general, functional enrichment highly varied by RD.

**Fig. S6.**
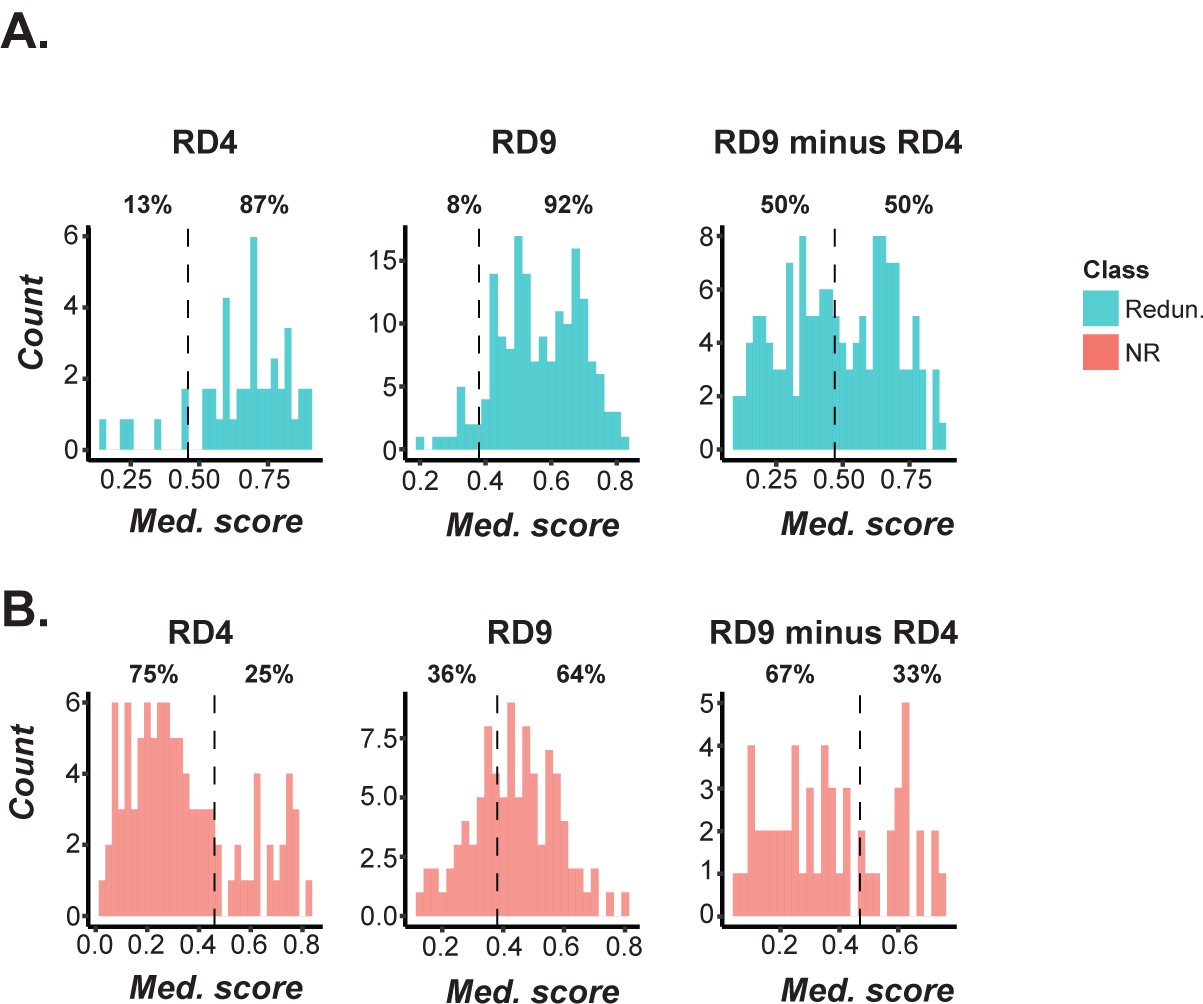
(*A-B*) Performance in cross-validation; the percentages of (*A*) redundant and (*B*) nonredundant gene pairs correctly and incorrectly predicted using different RDs are shown. Gene pairs to the left of the threshold (dotted line) were classified as nonredundant and gene pairs to the right were classified as redundant. Models were built using RD4 pairs, RD9 pairs, and RD9 pairs not included in RD4 as the redundant instances (RD9 minus RD4) with a randomly-selected half of nonredundant pairs as the nonredundant instances.

**Fig. S7.**
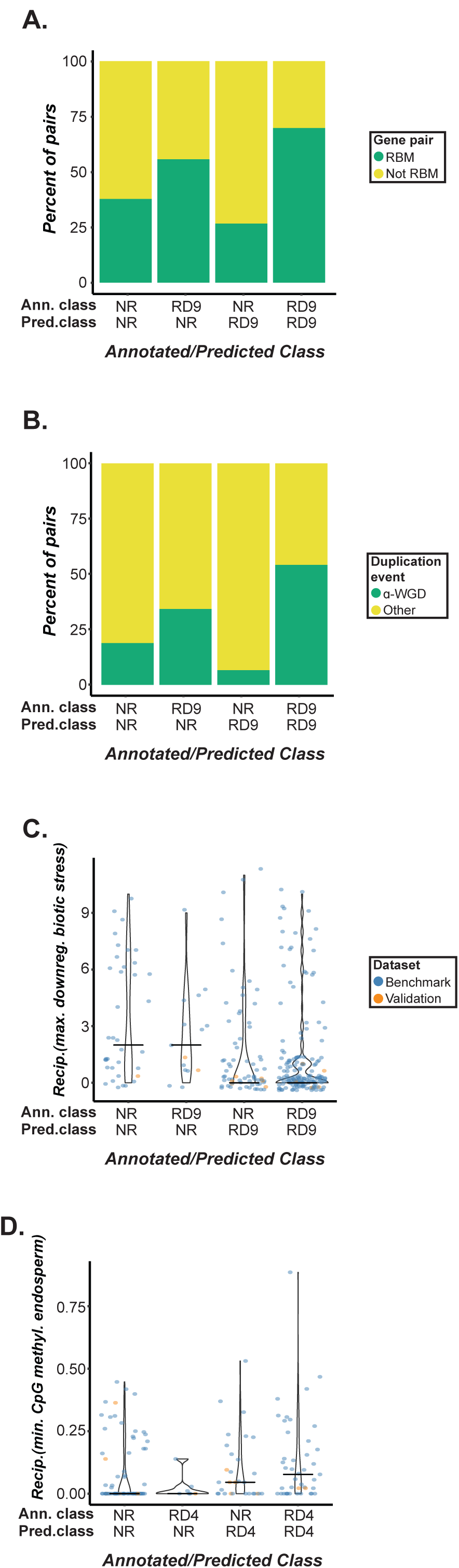
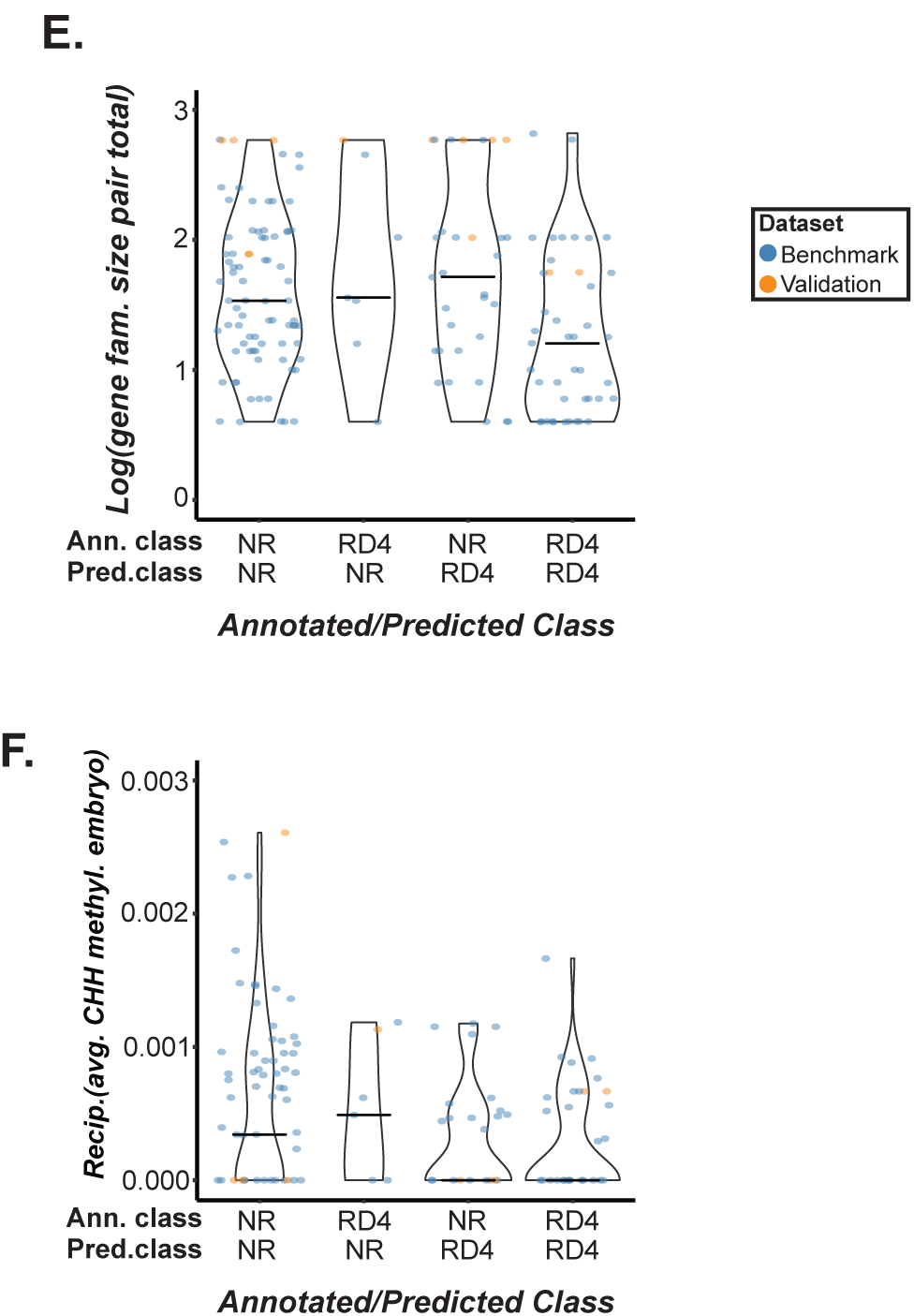
(*A*) Distribution of reciprocal best match gene pairs (RBM) among annotated (Ann.) vs. predicted (Pred.) RD9 classes: true negatives (NR/NR), false negatives (RD9/NR), false positives (NR/RD9), and true positives (RD9/RD9), including the benchmark dataset and the 10 validation pairs identified from the literature (here referred to as validation pairs). Two NR/RD9 validation pairs were reciprocal best matches, which was observed more often for RD9/RD9 pairs than NR/NR pairs, while genes in the RD9/NR validation pair were not reciprocal best matches, likely explaining these three mis-predictions. (*B*) Distribution of ɑ-whole genome duplication (ɑ-WGD)-derived gene pairs among the annotated/predicted classes, including the benchmark and 10 validation pairs. Three validation NR/RD9 pairs were derived from the ɑ-WGD event, which was observed more often for RD9/RD9 pairs than NR/NR pairs, potentially contributing to their mis-prediction. (*C*) Distribution of feature values among benchmark and validation gene pairs for the maximum biotic downregulation breadth between genes in a pair; a reciprocal transformation was applied to generate reciprocal maximum biotic downregulation breadth. All five of the validation NR/RD9 pairs had high values for this feature and looked more similar to RD9/RD9 pairs than to NR/NR pairs. (*D*) Distribution of reciprocal minimum CpG methylation in endosperm cells values among benchmark and validation pairs. (*E*) Distribution of total gene family size values among benchmark and validation pairs. The RD4/NR validation pair had a high value, which was more consistent with the values of NR/NR pairs than RD4/RD4 pairs. (*F*) Distribution of reciprocal average CHH methylation in embryo tissue values among benchmark and validation pairs. All four of the NR/RD4 validation pairs had high values that were more similar to those of RD4/RD4 gene pairs, while the one RD4/NR pair had a low value more similar to those of NR/NR pairs.

